# A combinatorial pre-mRNP retention factor network couples nuclear speckle architecture to splicing quality control

**DOI:** 10.64898/2026.09.28.754972

**Authors:** Kerstin Dörner, Kai Kurth, Nicole Beuret, Nikolaus Beer, Michael K. Jones, Karen Davey, Lea Stadelmann, Dominik Hügli, Alexia Ferrand, Giulia Basile, Seraphine Lüscher, Jernej Ule, Franka Voigt, Maria Hondele

## Abstract

Cells must prevent incompletely spliced transcripts from entering the cytoplasm, yet how splicing status is recognized and converted into selective nuclear retention remains unclear. Here, we identify a network of RNA-binding proteins that retain pre-mRNPs within nuclear speckles. Proteomic analysis of splicing intermediates accumulated in nuclear speckles identified a set of retention factors including BCLAF1, THRAP3 and SRSF7 that anchor pre-mRNPs to the SON scaffold and are required for pre-mRNP retention in speckles. Distinct RNA-binding patterns and transcript-specific retention factor requirements suggest that multiple mRNP features tune retention combinatorially. Super-resolution imaging revealed that the SON and SRRM2 speckle scaffolds form interdigitated networks with RNA enriched along networks and within interstitial speckle lumens. This architecture remodels in response to pre-mRNP load and is disrupted upon depletion of retention factors. Together, these findings define a pre-mRNP quality-control pathway that links splicing state to nuclear retention and the dynamic organization of nuclear speckles.

## Introduction

Most human protein-coding transcripts are synthesized as pre-mRNAs in which exons are interrupted by multiple introns. Introns are removed by the spliceosome, a dynamic ribonucleoprotein machinery built from snRNP particles that assemble stepwise on the pre-mRNA ^1^. Splicing, however, is neither instantaneous, nor uniformly efficient: weak splice sites and challenging intron architectures can delay splicing ^2–4^. This creates a fundamental sorting problem: incompletely processed transcripts must be retained in the nucleus long enough for splicing to complete, or for RNA surveillance to intervene, while their premature export must be prevented ^5–7^. The molecular basis of this retention and its spatial organization within the nucleus is however only beginning to emerge.

Nuclear speckles are emerging as important compartments at the interface between pre-mRNA processing and export competence. Organized around the scaffold proteins SON and SRRM2, they concentrate spliceosomal and pre-mRNP-maturation factors ^8–10^. Highly expressed, gene-rich chromosomal regions preferentially lie near nuclear speckles ^11^, bringing nascent transcripts into an environment that promotes spliceosome recruitment and co-transcriptional splicing ^12^. Speckles seem to be particularly important for GC-rich genes with short introns clustered within large, uniformly GC-rich genomic regions (GC isochores), whose processing is preferentially impaired when speckle organization is disrupted ^13^.

Yet nuclear speckles also appear to function beyond co-transcriptional splicing: intron-retaining transcripts are enriched at nuclear speckles ^14^, transcripts with inefficiently excised introns show more stable speckle enrichment ^15^, and experimentally perturbing spliceosome progression causes poly(A)+ RNA and spliceosomal components to accumulate in enlarged nuclear speckles ^16^. In addition, speckles can also transiently retain fully processed, export-competent mRNAs when transcription is inhibited ^17^, indicating that nuclear speckles can accommodate pre-mRNPs in distinct processing states.

How incompletely processed mRNPs are selectively recognized and retained within nuclear speckles, and which machinery mediates such retention, remains unclear. In reporter RNAs, an intact 5′ splice-site motif increases speckle association in a manner dependent on U1 snRNP proteins U1-70K and ZFC3H1 ^18^, and splice-site features that recruit U1 snRNP and U2AF can promote RNA localization to speckles ^19^. Moreover, the quality control factor LENG8 is recruited to intron-retaining pre-mRNAs by U1 snRNP and early splicing factors, preventing their nuclear export by antagonizing TREX-2 and promoting their degradation ^20^. Together, these studies show that cis-acting RNA features and engagement of early splicing factors can promote nuclear speckle localization or restrict export. What remains unknown is the trans-acting machinery that anchors these pre-mRNPs within nuclear speckles.

Here, we identify a network of RNA-binding proteins that associates with early spliceosomal pre-mRNPs and facilitates their retention within nuclear speckles, thereby counteracting premature nuclear export. Individual transcripts depend on overlapping but distinct components of this retention factor network, consistent with combinatorial recognition rather than a single universal retention signal. Retention factors connect pre-mRNPs to the SON scaffold and retained pre-mRNPs remodel the internal architecture of nuclear speckles. These findings define a selective retention pathway that anchors incompletely processed mRNPs within nuclear speckles and reveal retained pre-mRNPs as determinants of speckle organization.

## Results

### Arrest at early spliceosomal transitions routes polyA(+) mRNPs to nuclear speckles

How pre-mRNPs are recognized and prevented from prematurely acquiring export competence remains poorly understood. We reasoned that retention of incompletely spliced transcripts in speckles is mediated by RNA-binding proteins that anchor them to the nuclear speckle scaffold. To identify such factors, we looked for a means to enrich retained pre-mRNPs. In untreated cells only few pre-mRNPs are retained. Chemical splicing inhibitors efficiently amplify poly(A)+ RNA accumulation in nuclear speckles but provide no molecular handle on the underlying pre-mRNPs. We therefore exploited spliceosomal DEAD/H-box ATPases (DDXs/DHXs), which drive remodeling transitions throughout the splicing cycle (Figure S1A). ATPase-deficient DDX^DQAD^ and DHX^AQAD^ variants often arrest these transitions while remaining stably associated with their pre-mRNP substrates (Figure S1B), providing potential baits for isolating defined spliceosomal intermediates that engage the retention pathway ^21–23^.

To identify spliceosome perturbations that robustly induce nuclear speckle retention, we transiently expressed wild-type or ATPase-deficient variants of 16 DDX/DHX ATPases implicated in pre-mRNA splicing or nuclear speckle biology and monitored speckle morphology by SRRM2 immunofluorescence. Because accumulation of incompletely processed mRNPs is accompanied by nuclear speckle enlargement ^16,24^, we used increased speckle size as an initial phenotypic readout. DDX42^DQAD^, DDX23^DQAD^ and DHX16^AQAD^ produced the strongest nuclear speckle enlargement, whereas the remaining ATPases showed only modest or no detectable enlargement (Figures 1A-D, S1C and S2A). Thus, arrest at several early, pre-catalytic spliceosomal remodeling steps is particularly effective at driving nuclear speckle enlargement.

**Figure 1.**
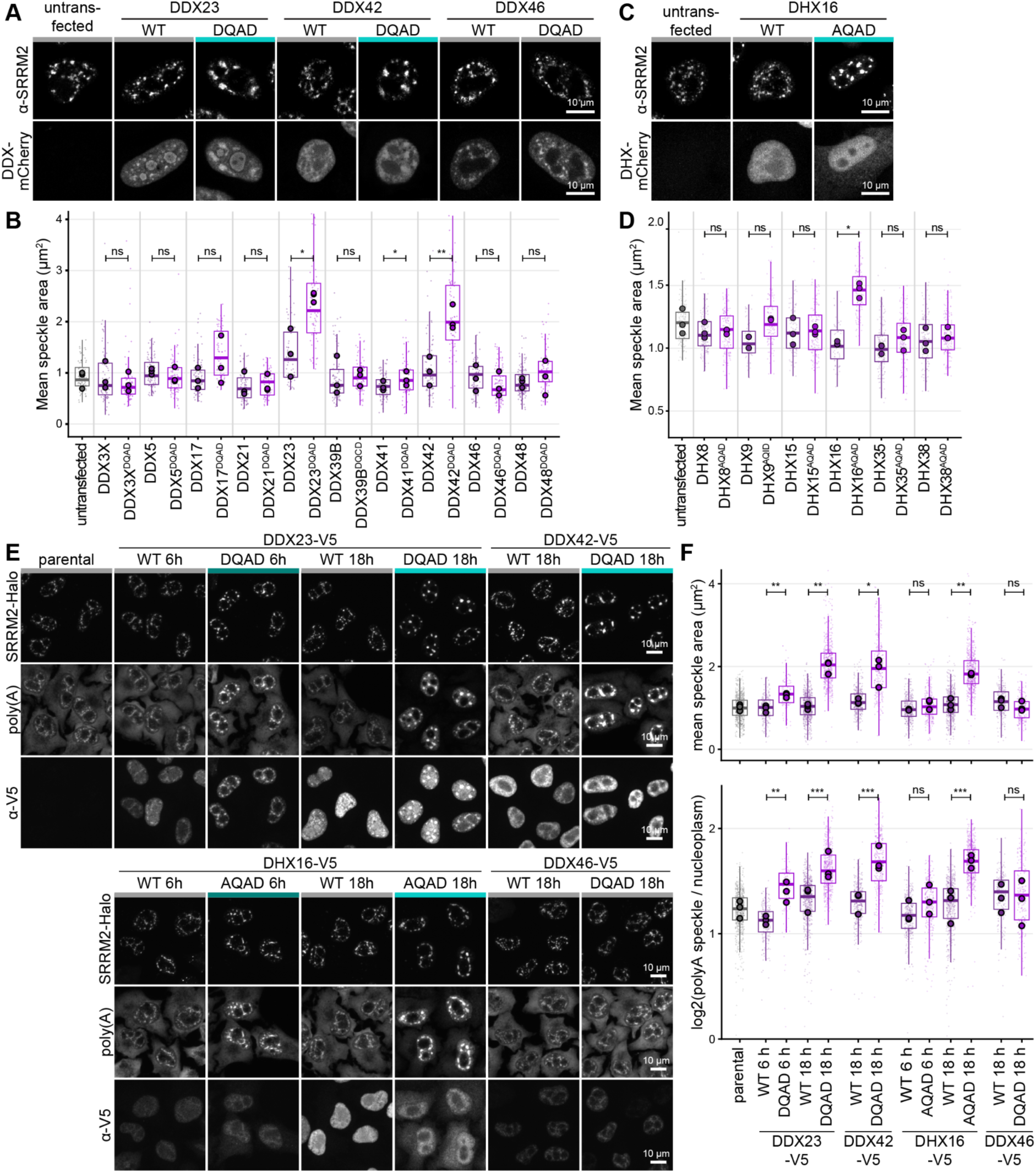
ATPase-deficient spliceosomal helicases drive poly(A)+ RNA accumulation in enlarged nuclear speckles. **(A)** Transient expression of C-terminally mCherry tagged DEAD-box (DDX) ATPase mutants, wild-type (WT) and ATPase-deficient (DQAD). 24 h after transfection cells were fixed and immunostained for SRRM2 using the SC35 antibody, imaged on . Mid nuclear planes for selected DDXs are shown; images of all tested DDXs are shown in Fig S1C. Turquoise color bars mark conditions with enlarged nuclear speckles. N = 3, n ≥ 60. Scale bars, 10 µm. **(B)** Quantification of nuclear speckle size in cells expressing the indicated WT or ATPase-deficient DDX variants in (A). For each cell, mean nuclear speckle area was calculated from SRRM2-defined speckles and converted to µm². Small points represent individual cells, boxes show the single-cell distribution, and large points represent the mean of each biological replicate. Dedicated untransfected control cells are shown in grey. Statistical significance was assessed using the three biological-replicate means by paired two-tailed t-tests comparing each DQAD mutant with the replicate-matched WT condition. p(DDX23) = 0.041, p(DDX41) = 0.036, p(DDX42) = 0.0035, *p < 0.05, **p < 0.01; ns, not significant. N=3. **(C)** Transient expression of C-terminally mCherry-tagged DEAH/DExH-box (DHX) ATPase mutants, wild-type (WT) and ATPase-deficient (AQAD). 24 h after transfection, cells were fixed and immunostained for SRRM2 using the SC35 antibody. Representative maximum-intensity projections of Z-stacks are shown for selected DHXs; results for all tested DHXs are shown in Fig S2A. Turquoise color bars mark conditions with enlarged nuclear speckles. N = 3 (except DHX9: N=2), n ≥ 68. Scale bars, 10 µm. **(D)** Quantification of cells transiently transfected with DHX-mCherry in (C). Cells were filtered for construct expression relative to the untransfected control. Nuclear speckle area per nucleus shown as boxplots. Small points represent individual cells, boxplots show the cell-level distributions, and large circles indicate the mean of each biological replicate. Statistical significance was assessed for matched WT–AQxD pairs using a two-tailed paired t-test on biological replicate means. p(DHX16 WT vs AQAD) = 0.0319 ; ns, not significant. **(E)** Stable SRRM2-Halo knockin HeLa cell lines were induced with doxycycline to express selected C-terminally V5-tagged DDX/DHXs mutants (WT or ATPase deficient DQAD/AQAD), at near-endogenous to moderately elevated levels for 18 h or 6 h as indicated. Cells were treated with TMR-Halo ligand for the last 6 h. After fixation, cells were stained for poly(A) RNA by FISH, followed by immunostaining against V5. Representative maximum-intensity Z-projections are shown. Turquoise color bars mark conditions with enlarged nuclear speckles. N=3, n≥ 122, Scale bars, 10 µm. **(F)** Quantification of induced V5-DDX/DHX cell lines in (E). Mean nuclear speckle area per nucleus and poly(A) RNA enrichment in nuclear speckles relative to the nucleoplasm are shown. Small points represent individual cells, boxplots show the cell-level distributions, and large circles indicate the mean of each biological replicate. Three biological replicates were analyzed. For nuclear speckle area statistical significance was assessed by two-tailed paired t-tests on replicate means of WT vs matched mutant p(DDX23 WT vs DQAD, 6h)= 0.0023; p(DDX23 WT vs DQAD, 18h)= 0.0011; p(DDX42 WT vs DQAD, 18h)= 0.0408; p(DHX16 WT vs AQAD, 18h)= 0.0096. For poly(A) localisation statistical significance was assessed by indicated matched WT–mutant comparisons using hierarchical linear mixed-effects models accounting for biological replicate, condition within replicate and imaging field, followed by planned contrasts with Benjamini–Hochberg correction across the six comparisons; p(DDX23 WT vs DQAD, 6h)= 0.0072; p(DDX23 WT vs DQAD, 18h)= 7.52×10^−4^; p(DDX42 WT vs DQAD, 18h)= 1.42×10^−4^; p(DHX16 WT vs AQAD, 18h)=1.35×10^−5^.

To validate these phenotypes under controlled expression conditions and determine whether speckle enlargement reflected RNA accumulation, we generated a homozygous SRRM2-Halo CRISPR knock-in HeLa line and, in this background, doxycycline-inducible cell lines carrying V5-tagged wild-type or ATPase-deficient DDX23, DDX42 and DHX16 constructs, and DDX46 as negative control (Figures S2B-E). Doxycycline amount and induction time was titrated to achieve expression levels comparable to the corresponding endogenous proteins (Figure S2F). As in the transient transfection screen, DDX23^DQAD^, DHX16^AQAD^, and DDX42^DQAD^ reproducibly enlarged nuclear speckles, whereas the wild-type proteins as well as DDX46^WT/DQAD^ had little effect (Figures 1E, 1F and S2G). These morphological changes were accompanied by a pronounced nuclear accumulation of polyadenylated RNA (poly(A)+ RNA), which was most prominent in nuclear speckles (Figure 1F and S2G). The phenotype was most consistently observed with DDX23^DQAD^, making it a suitable bait for isolating spliceosomal intermediates that engage the retention pathway.

### DDX23^DQAD^-induced retention requires incorporation of DDX23 into a stalled spliceosomal mRNP

DDX23 contains an extended serine/arginine (SR) rich intrinsically disordered region (IDR), raising the possibility that DDX23 itself – like other SR proteins – is targeted to nuclear speckles via its IDR via its IDR (Figure S3A). Serial truncation analysis narrowed the region required for the DDX23^DQAD^-induced phenotype to residues 162–165 within the N-terminal IDR (Figures S3A-C), and mutation of K162/F163 abolished the DQAD induced retention phenotype both after transient expression and in stable cell lines (Figures S3D, S3E and S4A– C).

Affinity purification-mass spectrometry (AP-MS) showed that disruption of the K162/F163 interface reduced association of DDX23 with mature spliceosomal assemblies while preserving association with U5 snRNP components and selectively enriching the U5 recycling factor TSSC4 (Figures S4D-F). Inspection of the human U4/U6.U5 tri-snRNP structure (PDB 6QW6 ^25^) revealed that DDX23 K162/F163 are positioned on the interface with the spliceosomal helicase SNRNP200, and structural comparison with spliceosome recycling complexes (PDBs 6QW6, 7PX3 ^25,26^) revealed that TSSC4 engages an overlapping surface on SNRNP200, with DDX23 F163 and TSSC4 F215 occupying the same hydrophobic pocket (Figures S4G and S4H) ^26^. These observations are consistent with disruption of the DDX23– SNRNP200 interface trapping DDX23 in an immature U5-associated state rather than allowing productive incorporation into the spliceosomal intermediate. Thus, the DDX23^DQAD^ retention phenotype requires formation of a stalled spliceosomal mRNP and is not explained by IDR-driven speckle-targeting activity of DDX23 itself, validating DDX23^DQAD^ as a bait for isolating speckle-retained spliceosomal complexes.

### Spliceosome-arrested pre-mRNPs reveal a candidate retention-factor network

If retention of pre-mRNPs in nuclear speckles is not encoded in the SR-rich region of DDX23, this information should instead reside in proteins that associate with retained pre-mRNP complexes.We therefore compared proteins associated with DDX23^WT^ and DDX23^DQAD^ by AP-MS. The two baits recovered distinct spliceosomal assemblies: DDX23^DQAD^ preferentially associated with early spliceosomal components, including U1-70K/SNRNP70, consistent with accumulation of a pre-B/pre-B-like intermediate in which U1 release has not occurred, whereas DDX23^WT^ recovered a broader, later spliceosomal composition (Figures 2A and S5A, S1A). Beyond these expected differences in spliceosomal composition, DDX23^DQAD^ enriched a distinct group of RNA-binding proteins. Most prominent were the SR-like splicing factors BCLAF1 and THRAP3, together with the SR proteins SRSF6, SRSF7, SRSF10, TRA2A and TRA2B. We therefore considered these seven proteins candidate components of a pre-mRNP-retention network, herein referred to as candidate retention factors.

**Figure 2:**
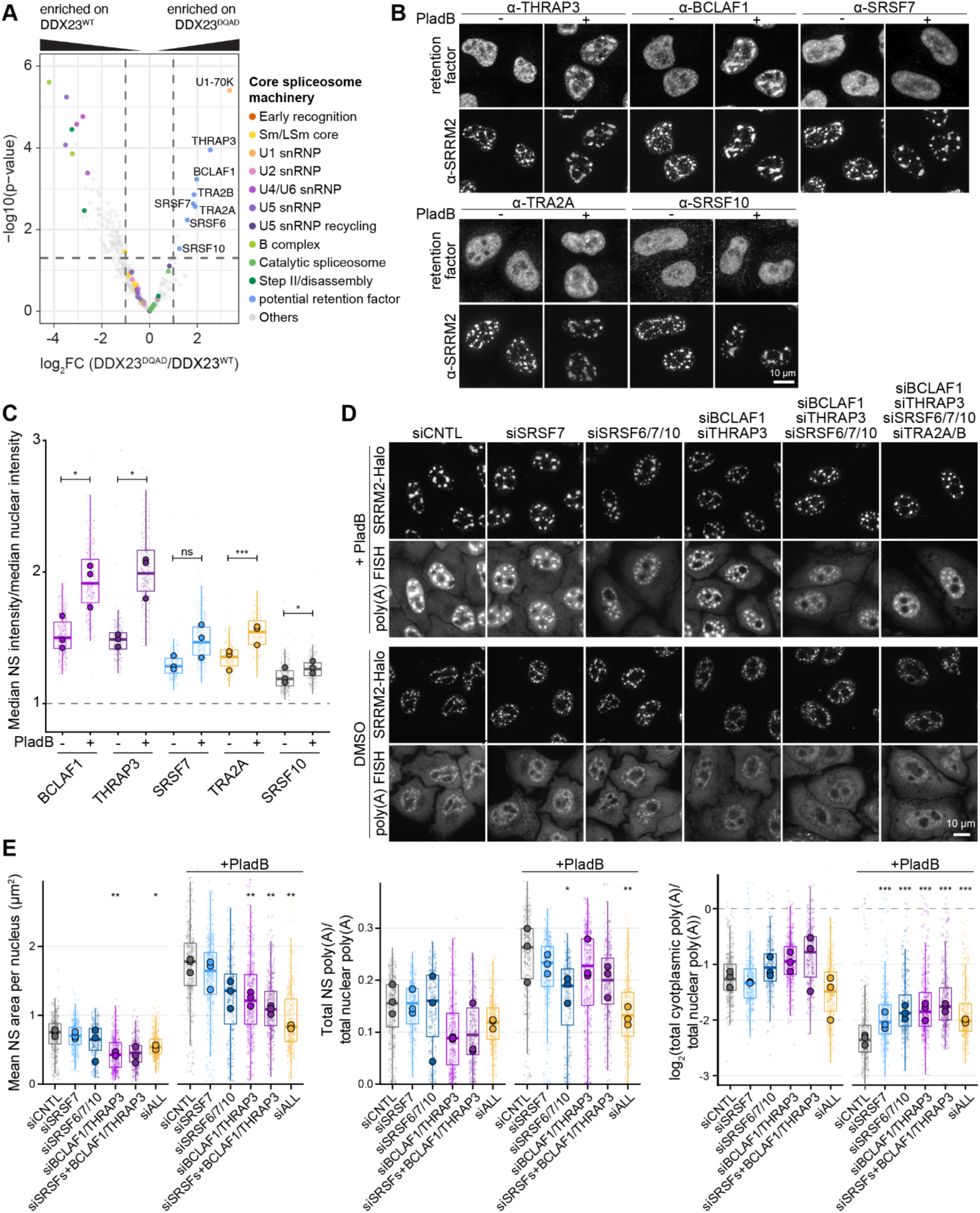
A network of RNA-binding proteins promotes poly(A)+ RNA retention in nuclear speckles. **(A)** Affinity purification–mass spectrometry (AP-MS) of DDX23, from the same experiment as Fig S4D. HEK293T cells expressing TwinStrep-HA–tagged DDX23 WT and DQAD were affinity-purified using Strep-Tactin beads, eluted with biotin and analyzed by mass spectrometry. The volcano plot shows enrichment versus significance; retention-factor candidates are labeled. N = 3. **(B)** HeLa Kyoto cells were either left untreated or treated with pladienolide B (PladB, 500 nM, 2.5 h), fixed, and immunostained with the indicated antibodies. Representative maximum-intensity projections are shown. N = 3, n ≥ 219. Scale bars, 10 µm. **(C)** Quantification of (B). For each cell, retention factor enrichment in nuclear speckles was calculated as the ratio of the median retention factor intensity within SRRM2-defined speckles to the median retention factor intensity across the nucleus. Box plots show the distribution across pooled single cells (line, median; box, interquartile range; whiskers, 1.5× IQR; outliers not shown); overlaid points are the means of the three biological replicates. A ratio of 1 (dashed line) indicates no speckle enrichment. Statistical significance was assessed by a two-tailed paired t-test on the biological-replicate means (N = 3; n ≥ 219 cells per condition): BCLAF1 p = 0.042, THRAP3 p = 0.020, TRA2A p = 1×10⁻⁴, SRSF10 p = 0.021, SRSF7 p = 0.22. ns, not significant. **(D)** HeLa SRRM2-Halo knockin cells were depleted of the indicated proteins by siRNA and SRRM2-Halo was labeled with TMR-Halo dye for 18 h. Cells were either left untreated or treated with pladienolide B (PladB, 500 nM, 2.5 h) before fixation. Poly(A) RNA was detected by FISH. Representative maximum-intensity projections are shown. N = 3, n ≥ 231. Scale bars, 10 µm. **(E)** Quantification of cells depleted of the indicated proteins in (D). Left, mean nuclear speckle area per nucleus. Middle, fraction of nuclear poly(A) signal contained within nuclear speckles, calculated for each cell as the summed integrated poly(A) intensity of all SRRM2-defined speckles divided by the integrated poly(A) intensity of the nucleus. Right, integrated poly(A) intensity cytoplasm / nuclear log₂-transformed. Box plots show pooled single-cell distributions (center line, median; box, interquartile range; whiskers, 1.5× IQR); large points indicate biological-replicate medians. For nuclear speckle area statistical significance was assessed using two-tailed t-tests on biological-replicate medians comparing each depletion with siCNTL within the corresponding treatment condition. For nuclear speckle area: p(siBCLAF1/THRAP3 vs siCNTL, noPladB)=0.0015; p(siSRSFs/BCLAF1/THRAP3 vs siCNTL, noPladB)=0.0015; p(siALLvs siCNTL, noPladB)=0.044; p(siBCLAF1/THRAP3 vs siCNTL, PladB)=0.035; p(siSRSFs+BCLAF1/THRAP3 vs siCNTL, PladB)=0.012; p(siALL vs siCNTL, PladB)=0.0028. For the fraction of nuclear poly(A) signal in speckles and nuclear/cytoplasm poly(A) ratio, statistical significance was assessed using a hierarchical linear mixed-effects model comparing each depletion with siCNTL within the corresponding treatment condition, with biological replicate and image incorporated as random effects; P values were corrected for multiple comparisons using the Benjamini–Hochberg method. For fraction of nuclear poly(A) in nuclear speckles: p(siSRSFs vs siCNTL, PladB)=0.014; p(siALL vs siCNTL, PladB)=0.0019. For ratio nuclear/cytoplasm poly(A) ratio: p(siSRSF7 vs siCNTL, PladB)=0.00037; p(siSRSFs vs siCNTL, PladB)=6.5×10^−8^; p(siBCLAF1/THRAP3 vs siCNTL, PladB)=2.5×10^−9^; p(siBCLAF1/THRAP3/SRSFs vs siCNTL, PladB)=7.0×10^−12^; p(siALL vs siCNTL, PladB)=3.4×10^−5^.

### A partially redundant RBP network promotes pre-mRNP retention in nuclear speckle

If these proteins are components of a pre-mRNP-retention machinery, increasing the abundance of incompletely processed mRNPs should increase their engagement with nuclear speckles. Under basal conditions, the candidate retention factors were distributed throughout the nucleoplasm, with only limited nuclear speckle enrichment (Figure S5B). Expression of DDX23^DQAD^ increased speckle enrichment of the candidate retention factors, most strongly BCLAF1 and THRAP3, consistent with their strong enrichment on DDX23^DQAD^-associated complexes (Figures 2A, S5C and S5D). Acute inhibition of splicing with the small molecule inhibitor Pladienolide B (PladB), which targets SF3B1 and thus an earlier step than DDX23^DQAD 27,28^, produced even stronger recruitment of the candidate retention factors to nuclear speckles, although the extent of redistribution differed between them (Figures 2B and 2C).

This redistribution was not simply a generic consequence of altered speckle morphology: inhibition of CLK kinases, which impairs splicing, SR-protein phosphorylation and SR-protein release from speckles ^29^, likewise promoted recruitment of several retention factors, with BCLAF1 and THRAP3 enrichment detectable before pronounced speckle enlargement (Figures S6A and S6C). By contrast, transcriptional inhibition with ActD or DRB altered speckle morphology, but did not induce changes in candidate retention factor localisation (Figures S6A–C). Because BCLAF1 and THRAP3 have been proposed to connect nuclear speckles to sites of active transcription ^30^, we also examined their localization relative to elongating RNA polymerase II (Ser2P). Their redistribution showed no consistent spatial relationship with Pol II foci, arguing that their redistribution to nuclear speckles is not simply driven by association with active transcription sites (Figure S6D).

Speckle recruitment of the candidate retention factors was partially interdependent, but the dependencies differed between perturbations. Following PladB treatment, SRSF7 depletion markedly reduced recruitment of THRAP3 and BCLAF1, whereas TRA2A/B depletion modestly increased THRAP3 accumulation (Figure S7A and S7B). Under CLK inhibition, this relationship was reversed: THRAP3 and BCLAF1 recruitment no longer depended on SRSF7, but was strongly reduced by TRA2A/B depletion (Figure S7A-C). TRA2A/B depletion also caused distinct THRAP3- and BCLAF1-positive foci outside nuclear speckles (Figure S7A-C). However, TRA2A/B depletion additionally causes pronounced mitotic defects, these dependencies cannot be interpreted as a simple recruitment hierarchy. Together, these contrasting dependencies indicate that the candidate factors form a dynamic, partially interdependent and context-dependent network. SRSF7 can facilitate THRAP3/BCLAF1 recruitment to speckles upon spliceosome inhibition, whereas altered SR-protein phosphorylation shifts this dependency toward TRA2A/B.

If these proteins form a pre-mRNP-retention network, their depletion should perturb pre-mRNP organization. Depletion of several retention factors in combination indeed altered nuclear speckle morphology and poly(A)+ RNA accumulation in untreated cells (Figures 2D, 2E and S8). When we increased the load of incompletely processed pre-mRNPs with PladB, depletion of SRSF7, BCLAF1/THRAP3 or combinations of retention factors reduced PladB-induced speckle enlargement and poly(A)+ RNA accumulation, with the strongest effects observed after combined depletion of all seven factors (Figures 2D, 2E and S8). The stronger phenotypes produced by combined depletion support partially overlapping contributions.

Depletion of SRSF7, BCLAF1/THRAP3 or all seven retention factors not only reduced nuclear speckle accumulation, but also lowered the nuclear-to-cytoplasmic poly(A)+ RNA ratio, most prominently following PladB treatment (Figure 2E). Thus, nuclear speckle retention is linked to the broader nuclear confinement of poly(A)+ pre-mRNPs.

Together, these findings establish SRSF7, BCLAF1, THRAP3 and the associated SR proteins as components of a functional pre-mRNP-retention pathway (‘retention factors’) that promotes sequestration within nuclear speckles and nuclear confinement. Nuclear speckles therefore emerge not simply as sites of pre-mRNP accumulation, but as quality-control compartments that help prevent incompletely processed transcripts from entering the cytoplasmic mRNP pool before processing is complete.

### Individual transcripts display distinct retention-factor dependencies

The partial phenotypes produced by depletion of individual retention factors suggested that different incompletely processed mRNPs may rely on distinct components of the retention network. To address this possibility, we selected *BRD4* and *KPNA1*, two endogenous transcripts previously reported to accumulate in nuclear speckles following transcription or transcription or spliceosome inhibition ^15,17,31^, and monitored their localization using single-molecule RNA FISH (smFISH). BRD4 was detected using exonic probes, whereas KPNA1 was analyzed with independent exonic and intronic probe sets.

Individual single transcripts of both BRD4 and KPNA1 localize to nuclear speckles in control cells, and retention factor depletion decreased this association, indicating that the retention pathway is important for their proper distribution in the nucleus (Figure S9A-C). PladB further increased the speckle enrichment of both BRD4 and KPNA1, providing a larger dynamic range in which to resolve their factor dependencies (Figures 3A-D). The two transcripts displayed overlapping but distinct retention factor requirements. BRD4 accumulation was reduced by SRSF7 depletion, and simultaneous depletion of all seven retention factors (siALL) largely abolished the PladB-induced increase in speckle enrichment (Figures 3A and 3B). KPNA1 depended on both SRSF7 and BCLAF1/THRAP3: depletion of either reduced speckle enrichment of both the exonic and intronic KPNA1 signals, linking these dependencies directly to incompletely spliced KPNA1 transcripts. Combined depletion of all seven factors further reduced KPNA1 accumulation (Figures 3C and 3D).

**Figure 3:**
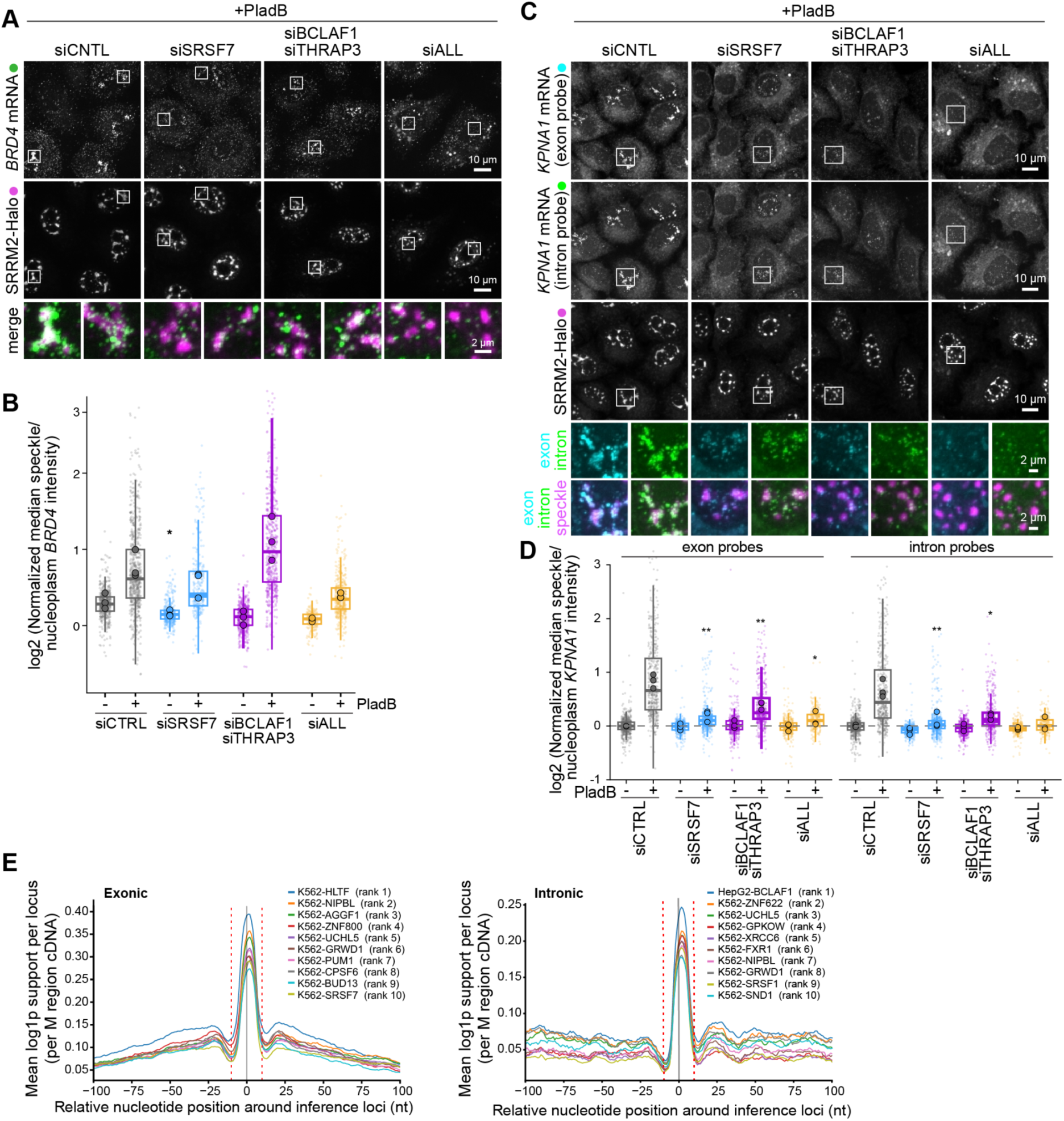
Different mRNAs depend on distinct combinations of retention factors for nuclear speckle accumulation. **(A)** HeLa SRRM2-Halo knockin cells were depleted of the indicated factors by siRNA, cells were treated with Janelia Fluor 503 HaloTag ligand for 18 h. Cells were either treated with PladB (500 nM, 2.5 h). After fixation single mRNAs were stained with probes targeting exonic sequences of *BRD4* mRNA using smFISH. Representative maximum-intensity projections are shown. N = 3, n ≥ 209. Scale bars, 10 µm, enlargements, 2 µm. **(B)** Quantification of (A). Background-corrected median BRD4 speckle/nucleoplasm intensity ratio per cell, normalized within each replicate to siCTRL −PladB and log₂-transformed (0 = control, dashed line). Boxes, median and IQR of single cells (faint points); large circles, per-replicate means (N = 3 biological replicates). Significance was tested on replicate means by two-tailed paired t-test versus the matched siCTRL, p(siSRSF7 -PladB vs siCNTL -PladB) =0.048. PladB increased enrichment 1.8-fold in control cells (p = 0.014). **(C)** HeLa SRRM2-Halo knockin cells were depleted of the indicated factors by siRNA, cells were treated with Janelia Fluor 503 HaloTag ligand for 18h. Cells were either treated with PladB (500 nM, 2.5h). Cells left untreated are shown in Figure S9. After fixation, single mRNAs were stained with a pool of probes targeting either intronic or exonic sequences of *KPNA1* mRNA using smFISH. Representative maximum-intensity projections are shown. N = 3, n ≥230. Scale bars, 10 µm, enlargements, 2 µm. **(D)** Quantification of (C). Background-corrected median *KPNA1* speckle/nucleoplasm intensity ratio per cell, normalized within each replicate to siCTRL −PladB and log₂-transformed (0 = control, dashed line). Boxes, median and IQR of single cells (faint points); large circles, per-replicate means (N = 3 biological replicates). Significance was tested on replicate means by two-tailed paired t-test versus the matched siCTRL, p(exon: siSRSF7 +PladB vs siCNTL +PladB) =0.0014, p(exon: siBCLAF1/THRAP3 +PladB vs siCNTL +PladB) =0.0085, p(exon: siALL +PladB vs siCNTL +PladB) =0.04, p(intron: siSRSF7 +PladB vs siCNTL +PladB) =0.0026, p(intron: siBCLAF1/THRAP3 +PladB vs siCNTL +PladB) =0.028, p(intron: siALL +PladB vs siCNTL +PladB) =0.051. **(E)** RBPeek metaprofiles of ENCODE eCLIP co-binding around HA-THRAP3 iCLIP crosslink sites in HEK293T cells expressing HA-THRAP3 at near-endogenous levels (48 h induction). THRAP3 inference loci (single-nucleotide midpoints present in ≥2 of 4 replicates) were split into exonic (left, n = 19,917) and intronic (right, n = 8,666) sets. The top 10 of 224 datasets, ranked by central binding (area under the metaprofile within ±10 nt, red dotted lines; grey line = THRAP3 locus), are shown.

Individual transcripts thus seem to depend on distinct combinations of retention factors rather than a single universal retention component. The different requirements of BRD4 and KPNA1 further suggest that individual components of the network may recognize and integrate distinct features of incompletely processed mRNPs.

### Distinct RNA-binding modes separate functional modules of the retention network

The transcript-specific factor requirements suggested that the retention network may comprise functionally distinct RNA-binding components rather than a single shared recognition module. To examine this possibility, we profiled the RNA binding of THRAP3 by iCLIP in HEK293 cells, using anti-HA antibody against stably integrated HA-THRAP3. THRAP3 was found to bind predominantly to intronic and intergenic regions, with additional binding in coding sequences within exons (Figure S9D). Motif analysis by PEKA identified U-rich 5-mers as the top-enriched sequences in both introns and exons, and in addition GA-rich motifs in introns and A/AG-rich motifs in exons (Figure S9E and S9F).

To test whether THRAP3 shares its binding sites with other RNA binding proteins we compared THRAP3 crosslink sites with binding profiles of all 223 datasets of the ENCODE eCLIP compendium (150 RBPs; 103 and 120 RBPs in HepG2 and K562 cells, respectively) ^32,33^ and ranked them by their enrichment at exonic and intronic THRAP3 peaks (Figure 3E).

BCLAF1 showed the strongest enrichment of all proteins at intronic THRAP3 peaks, but did not rank highly among proteins overlapping at exonic peaks. SR/SR-like proteins of the retention factor network did not show much overlap, with only SRSF7 ranked ninth among the proteins that were enriched at the exonic crosslink peaks of THRAP3. Thus, THRAP3 and BCLAF1 occupy a closely shared RNA-binding environment in introns, while the SR-protein components of the retention network show some overlap to its exonic binding.These data indicate that the retention network engages incompletely processed mRNPs through distinct RNA-binding modules rather than a single shared recognition mode.

### Retention factors engage the nuclear speckle scaffold

Retention of incompletely processed mRNPs ultimately requires their association with the nuclear speckle scaffold. To determine how the retention machinery engages this environment, we mapped the proximity proteomes of SRSF7 and THRAP3 by APEX2-mediated proximity labeling followed by quantitative mass spectrometry in untreated cells and after PladB treatment (Figure S10A-E).

SRSF7 displayed a largely stable proximity landscape. In both untreated and PladB-treated conditions, it was proximal to other retention factors, spliceosomal proteins, and the nuclear speckle scaffold components SON and SRRM2 (Figures S10A, Figure 4A), consistent with a constitutive association with the speckle environment and the comparatively modest redistribution of SRSF7 following splicing inhibition (Figure 2B and 2C). THRAP3 behaved differently. Under basal conditions, THRAP3 was proximal to its known interaction partners BCLAF1 and ERH ^34^, as well as to other retention factors and spliceosomal proteins (Figures S10B, Figure 4A). As incompletely processed mRNPs accumulated following PladB treatment, its proximity to spliceosomal proteins as well as SON and SRRM2 increased. These data place both SRSF7 and THRAP3 within the nuclear speckle scaffold environment and show that THRAP3’s engagement with this environment increases with the load of incompletely processed mRNPs.

**Figure 4:**
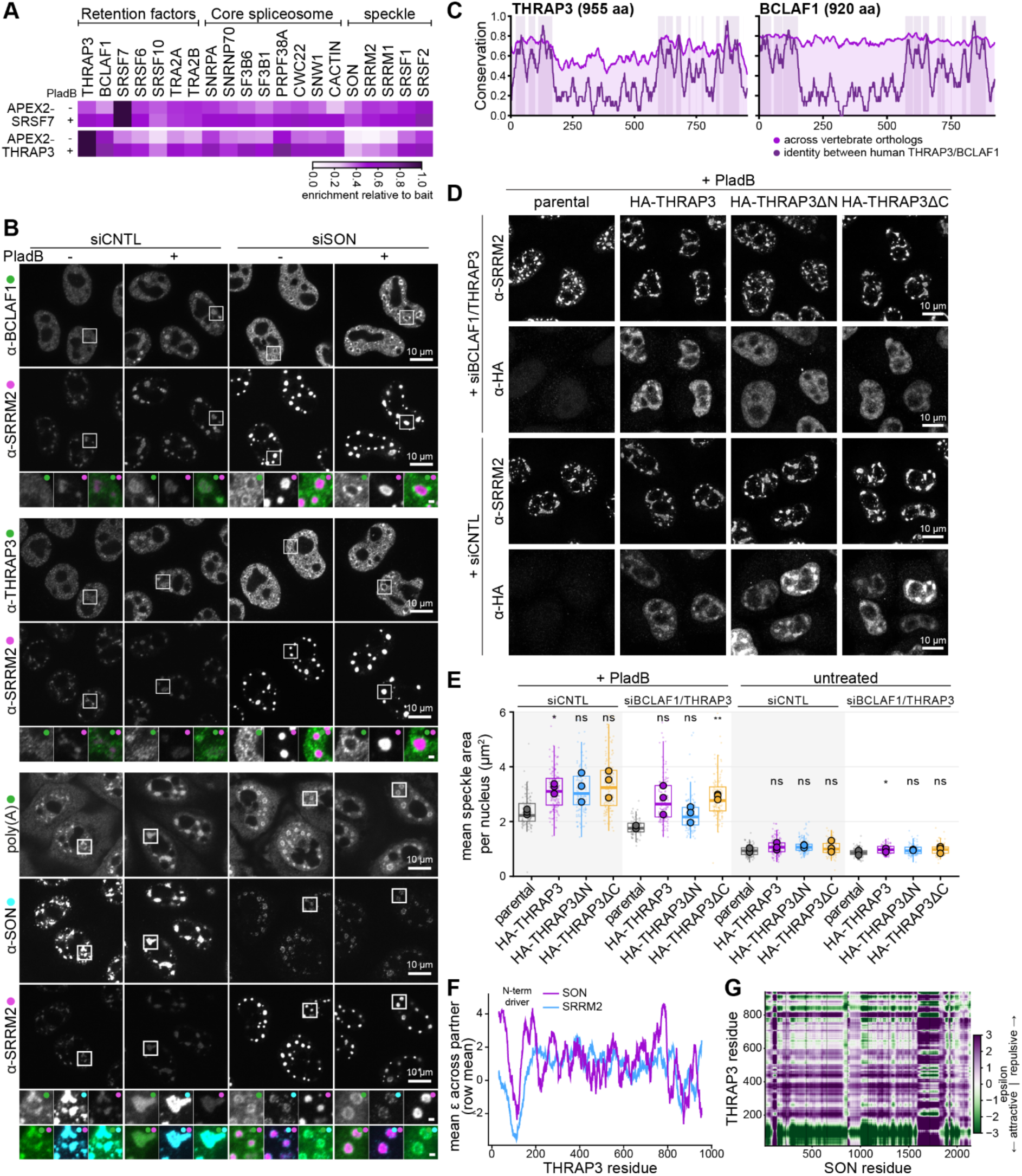
SON anchors retained pre-mRNPs to the nuclear speckle scaffold. **(A)** Heat map of APEX2 proximity-labeling proteomics for SRSF7 and THRAP3. Stable HeLa cell lines were induced to express APEX2-V5-SRSF7 or APEX2-V5-THRAP3 at near-endogenous levels for 48 h and either left untreated or treated with pladienolide B (PladB, 500 nM, 2.5 h) before APEX2-mediated proximity labeling and harvest; cells expressing APEX2-V5-NLS served as a control. Each cell shows enrichment of the indicated factor relative to the bait’s own self-enrichment in that sample (bait self-labeling = 1), so the two experiments share a common scale. N = 3. **(B)** HeLa cells were depleted of SON by siRNA (48 h). Cells were then treated with DMSO or PladB (500 nM) for 2.5 h before fixation. Poly(A) RNA was stained using FISH, proteins were then immunostained with the indicated antibodies. Representative single planes are shown. N = 3. Scale bars, 10 µm; enlargement 1 µm. **(C)** Sequence conservation and paralog-shared identity per-residue profiles for human THRAP3 and BCLAF1. Pink line shows ortholog conservation (Jensen–Shannon divergence relative to a BLOSUM62 background, computed from 188 THRAP3 and 132 BCLAF1 vertebrate orthologs and smoothed over a 15-residue window), and the violett line shows paralog-shared identity (fraction of a 21-residue window in which the THRAP3 and BCLAF1 ortholog consensus residues are identical). Violett shaded bands mark contiguous paralog-shared blocks (smoothed identity > 0.5). **(D)** Stable HeLa cell lines were induced to express HA-THRAP3, HA-THRAP3ΔN (HA-THRAP3^176–955^) or HA-THRAP3ΔC (HA-THRAP3>^1–828^) at near-endogenous protein level. Cells were then either co-depleted for BCLAF1 and THRAP3 by siRNA or treated with control siRNA. Cells were treated with PladB (500 nM, 2.5 h) before fixation and immunostained with the indicated antibodies. Representative maximum-intensity projections are shown. N = 3, n ≥ 80. Scale bars, 10 µm. **(E)** Quantification of stable cell lines co-depleted of BCLAF1 and THRAP3 in (D) and Figure S11C. Box plots show the mean nuclear speckle area per cell (line, median; box, interquartile range; whiskers, 1.5× IQR); overlaid points are the means of the three biological replicates. Statistical significance was assessed by a two-tailed paired t-test on the biological-replicate means (N = 3; n ≥ 80 cells per condition): p(HA-THRAP3 vs parental, +PladB, siCNTL)=0.029, p(HA-THRAP3 vs parental, untreated, siBCLAF1/THRAP3)=0.029, p(HA-THRAP3ΔC vs parental, +PladB, siBCLAF1/THRAP3)=0.0065, ; ns, not significant. **(F)** Per-residue interaction profile of full-length THRAP3 against SON (purple) and SRRM2 (blue), predicted with FINCHES (Mpipi frontend). Each point is the mean-field interaction parameter ε for the sequence window centered on that THRAP3 residue, averaged across the entire partner protein (row-mean of the maps in Fig 4G; 31-residue sliding window); ε < 0 indicates net attraction and ε > 0 net repulsion. **(G)** Residue-resolved FINCHES interaction maps for THRAP3 × SON. Each pixel is the predicted ε between the local windows centered on the corresponding THRAP3 residue (y-axis) and partner residue (x-axis); green denotes attraction (ε < 0) and purple repulsion (ε > 0), on the shared scale at right (ε ∈ [−3, 3]). SON was tiled into non-overlapping ∼250-residue fragments (boundaries placed outside predicted folded islands), each run against full-length THRAP3 and stitched into the full-length maps shown.

### SON anchors retained pre-mRNPs within nuclear speckles

The proximity of THRAP3 and SRSF7 to SON and SRRM2 suggested that one or both major scaffold proteins could mediate the ‘anchoring’ of retained pre-mRNPs within nuclear speckles (Figure 4A). We therefore depleted SON or SRRM2 and examined the spatial organization of BCLAF1, THRAP3 and poly(A)+ RNA (Figure 4B and S11A). Depletion of either scaffold perturbed the organization of the remaining scaffold component and is known to impair splicing ^13^, complicating interpretation of changes in total RNA accumulation. We therefore focused on the spatial relationship between retained pre-mRNP components and the reorganized scaffold.

SRRM2 depletion reorganized SON into slightly smaller and brighter structures but had little effect on the accumulation of BCLAF1, THRAP3 or poly(A)+ RNA in the remaining nuclear speckle compartment after PladB treatment (Figure S11A). SON depletion produced a strikingly different organization (Figure 4B). SRRM2 collapsed into small, compact cores, with residual SON enriched in a peripheral domain surrounding these cores. BCLAF1, THRAP3 and poly(A)+ RNA were excluded from the SRRM2 cores and instead accumulated in the SON-containing peripheral domain in both untreated and PladB treated cells (Figure 4B and S11B). Line-profile analysis confirmed the spatial separation of BCLAF1, THRAP3 and poly(A)+ RNA from the SRRM2-rich core and their overlap with the peripheral SON-enriched region (Figure S11B). These observations identify SON as the principal nuclear speckle scaffold associated with retained pre-mRNPs.

To define how the retention machinery engages this scaffold, we mapped the region of THRAP3 required for retention. Sequence comparison across vertebrates revealed strong conservation and similarity of the N- and C-terminal regions of THRAP3 and BCLAF1 (Figure 4C). We therefore co-depleted endogenous BCLAF1 and THRAP3 and tested whether inducibly expressed full-length THRAP3 or variants lacking the N-terminal 175 residues (THRAP3ΔN) or C-terminal 127 residues (THRAP3ΔC) could rescue the nuclear speckle response.

Full-length THRAP3 and THRAP3ΔC restored speckle enlargement following splicing inhibition, whereas THRAP3ΔN failed to rescue (Figures 4D, 4E, S11C and S11D), identifying the conserved N-terminal region as essential for THRAP3-mediated pre-mRNP retention. FINCHES analysis ^35^ further predicted strong attractive interaction propensity between the THRAP3 N-terminus and SON, and weaker predicted interactions with SRRM2 (Figures 4F and 4G). Together with the preferential segregation of THRAP3 with SON, these findings suggest that interaction of the conserved THRAP3 N-terminus with SON contributes to retaining mRNPs at the nuclear speckle scaffold.

### The pre-mRNP retention network shapes nuclear speckles architecture

Anchoring of retained pre-mRNPs to the SON scaffold raised the possibility that the retention pathway contributes to the internal organization of nuclear speckles. From recent super-resolution studies, the emerging picture is that nuclear speckles are not homogeneous condensates but internally structured ‘patchy’ assemblies ^36–38^. Similar to this model, three-dimensional structured illumination microscopy (3D-SIM), revealed distinct SON and SRRM2 networks (Figure 5A and 5B), which, as judged by calculation of the occupancy matched overlap, are interdigitated and not superimposable. The networks are interspersed with internal regions of comparatively low scaffold density, which we refer to as ‘speckle lumens’. This organization was reproduced with reciprocal fluorophore labeling and by tauSTED imaging (Figures S12A-D).

**Figure 5:**
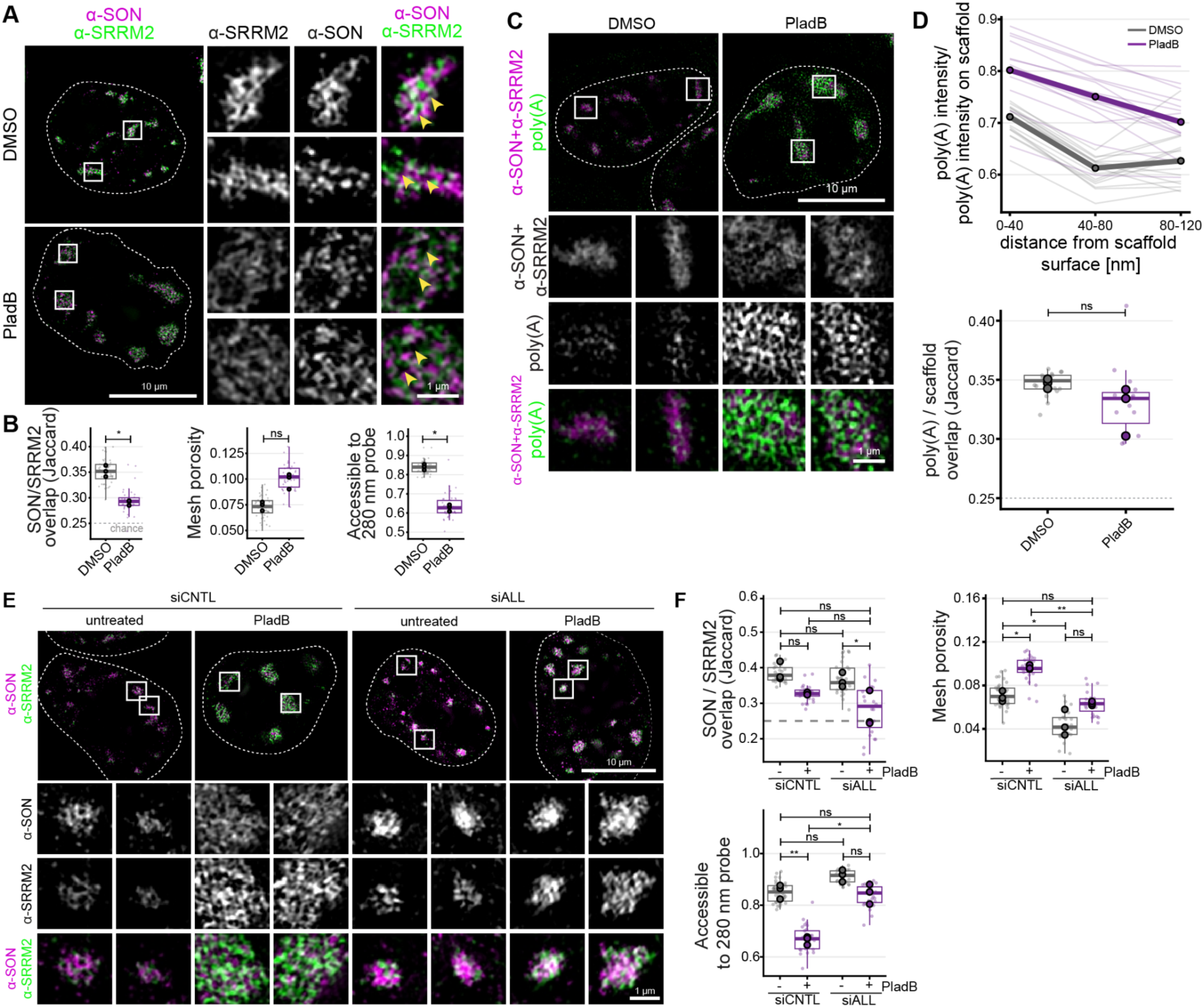
The pre-mRNP retention network shapes nuclear speckle architecture. **(A)** HeLa cells were treated with pladienolide B (500 nM, 2.5 h) or DMSO. After fixation, cells were immunostained with anti-SON labelled with a secondary antibody conjugated with Alexa488 and anti-SRRM2 labelled with a secondary antibody conjugated with Alexa405. Reciprocal stainings with swapped secondary colors are displayed in Figure S12A. Cells were imaged using 3D-SIM. Representative reconstructed single planes are shown. Yellow arrows highlight scaffold-poor regions. N = 3, n ≥ 41. Scale bars, 10 µm, enlargements 1 µm. **(B)** Quantification of cells in (A). Left, overlap of the SON- and SRRM2-dense phases within speckle bodies, measured as an occupancy-matched Jaccard index (top 40% voxels); the dashed line marks the chance level of 0.25, obtained when one channel is rotated 90 degrees within the same nucleus. Middle, mesh porosity of the resolved speckle core (fraction of core volume not occupied by SON/SRRM2 scaffold). Right, fraction of the intra- speckle interstitium reachable by a 280 nm spherical probe entering from the nucleoplasm. Small points, one value per cell (median over that cell’s speckles); large points, per-replicate medians; boxes, cell-level median and interquartile range with whiskers to 1.5x IQR. Statistics, paired two-sided t-test on the three replicate medians with Holm correction across the three panels: ns p > 0.05, * p < 0.05. p(SON/SRRM2 overlap DMSO vs PladB)=0.04; p(meshwork porosity DMSO vs PladB)=0.06334; p(interstitium accessible to 280 nm probe DMSO vs PladB)=0.02085 **(C)** Hela cells were treated with pladienolide B (500 nM, 2.5 h) or DMSO before fixation. Poly(A) mRNA was visualized using FISH, followed by staining with SON and SRRM2, both primary antibodies were stained using the secondary antibodies coupled to Alexa-405 fluorophores. Cells were imaged using 3D-SIM. Representative reconstructed single planes are shown. N = 3, n ≥ 15. Scale bars, 10 µm, enlargements 1 µm. **(D)** Quantification of cells in (C). Top: poly(A) intensity inside speckle bodies as a function of distance from the scaffold surface, expressed relative to the poly(A) intensity on the scaffold of the same cell, so the profile is independent of staining and exposure. Distance bins are 0– 40, 40–80, 80–120. Bins near 40–80 nm include a larger share of the sparse 80 nm boundary layer; the profile is monotonic when that layer is excluded. Thin lines, individual cells; heavy lines, condition medians. Bottom: occupancy-matched overlap of poly(A) with the scaffold (Jaccard of the top 40% of body voxels in each channel; dashed line, chance level 0.25, confirmed empirically by rotating the poly(A) channel 90° in xy: 0.252 DMSO, 0.258 PladB). Points, individual cells; large circles, replicate medians; boxes, median and interquartile range. Statistics: paired two-sided t-test on replicate medians, Holm-corrected across the panels shown; ns, p > 0.05. **(E)** Hela cells were co-depleted of THRAP3, BCLAF1, SRSF6/7/10, TRA2A/B by siRNA or treated with control siRNA. Before fixation, cells were treated with pladienolide B (500 nM, 2.5 h) or left untreated. After immunostaining with the indicated antibodies, cells were imaged using SIM. Representative reconstructed single planes are shown. N = 3, n ≥ 23. Scale bars, 10 µm, enlargements 1 µm. **(F)** Quantification of cells in (E) SON/SRRM2 overlap (Jaccard of the two channels thresholded at matched occupancy, 40% of body volume each; dashed line, chance = 0.25); mesh porosity of the resolved speckle core (fraction of core volume not occupied by SON/SRRM2 scaffold); fraction of the intra-speckle interstitium reachable by a 280 nm spherical probe entering from the nucleoplasm. Small points, individual cells; boxplots, cell- level distributions; large circles, median of each biological replicate. Two-tailed paired t-test on the three replicate medians, uncorrected. *p < 0.05, **p < 0.01, ns = not significant.

Poly(A)+ RNA was concentrated along the speckle scaffold but also extended into the surrounding lumens, with signal intensity gradually decreasing with distance from the scaffold (Figure 5C and 5D). Scaffold proximity, however, did not fully explain RNA organization. When calculating occupancy overlap, the spatial association of poly(A)+ RNA with the scaffold was comparable to that observed between the two scaffold proteins themselves, suggesting that additional factors contribute to the positioning of RNA within nuclear speckles. The same spatial organization was reproduced in independent SON/poly(A)+ RNA and SRRM2/poly(A)+ RNA 3D-SIM datasets and by Airyscan imaging (Figures S12E-G).

If speckle scaffold and lumens accommodate retained pre-mRNPs, their organization should respond to changes in RNA flux/content. Consistent with this prediction, splicing inhibition and the accompanying accumulation of poly(A)+ RNA drove spatial decompaction of the SON and SRRM2 scaffold networks, reduced their overlap and increased meshwork porosity, thereby increasing the volume of the lumens (Figure 5A, S12A-D). Poly(A)+ RNA signal remained overlapping with the scaffold but extended further into the enlarged lumens upon PladB treatment (Figure 5D and Figure S12G). Together, these findings indicate that retained pre-mRNPs are not passive cargo but contribute to remodeling the internal architecture of nuclear speckles.

To determine whether this RNA-responsive architecture depends on the retention machinery, we co-depleted all seven retention factors and examined SON/SRRM2 organization under basal conditions and following splicing inhibition. Loss of the retention network spatially compacted the characteristic SON/SRRM2 meshwork, reducing mesh porosity and thus lumen volume in both untreated and PladB-treated cells (Figures 5E, 5F, S12H and S12I). This failure to undergo scaffold decompaction upon retention-factor depletion likely provides the ultrastructural basis for the inability of these cells to enlarge nuclear speckles upon splicing inhibition (Figures 2D and S8). Thus, the retention pathway is important both for basal speckle architecture and for its adaptation to changes in pre-mRNP load.

Together, these findings show that pre-mRNP retention actively shapes nuclear speckle organization. By coupling incompletely processed mRNPs to the scaffold, the retention pathway remodels the internal meshwork and generates an RNA-responsive architecture whose network ultrastructure and luminal spaces expand and contract with changes in RNA load. Nuclear speckles therefore emerge as dynamic compartments whose architecture is continuously shaped by the flux and retention of likely incompletely processed pre-mRNPs.

## Discussion

Our findings support a model in which mRNP retention within nuclear speckles constitutes a regulated layer of quality control. Slow or stalled splicing may amplify this response, but the requirement for a defined RBP network indicates that nuclear and nuclear speckle confinement is not simply a passive consequence of delayed processing.

An important question is which features of an incompletely processed mRNP are recognized by this quality-control step. Previous work has linked intact 5′ splice-site signals and engagement of early splicing factors to speckle localization or nuclear retention ^18–20^. Our DDX screen complements these observations by showing that perturbations at distinct early stages of spliceosome assembly and remodeling induce nuclear speckle enlargement and poly(A)+ RNA accumulation. Several findings point to U1 snRNP-containing pre-mRNPs as retention-competent state: U1-70K is strongly enriched in DDX23^DQAD^-associated complexes, consistent with impaired U1 release at the DDX23-dependent transition, and U1-70K depletion reduces reporter 5’ splice site accumulation within nuclear speckles ^18^. This connection fits the recently proposed U1 relay model, which proposes that sequential U1-dependent interactions coordinate co-transcriptional splicing events and 3′-end formation ^39^.

A U1-associated splicing state alone, however, is unlikely to be the only determinant to explain why different mRNPs are retained with different efficiencies. Our results identify an additional layer of quality control: a network of RNA-binding proteins that associate with pre-mRNPs and are required for poly(A)+ RNA retention within nuclear speckles. This provides cells flexibility, allowing transcript-specific features to be read and retention to be tuned accordingly. These RBPs are known to recognise specific exonic and intronic sequence features to selectively regulate individual transcripts, suggesting that at least some speckle retention determinants result from combinatorial sequence-dependent recognition. Retention factors may also act beyond physiological pre-mRNP retention: during interstasis, TRA2 proteins promote the selective capture of purine-rich mRNAs within nuclear speckles, providing an example of sequence-dependent retention in a homeostatic response ^31^. Whether other factors identified here contribute to this response remains to be tested.

Retention also raises a spatial question: how are these mRNPs accommodated within the nuclear-speckle scaffold? Building on previous observations of internally segregated speckle substructures ^36,38^, our super-resolution imaging resolves SON and SRRM2 into a sponge-like scaffold surrounding poly(A)+ RNA-rich luminal spaces. When scaffold balance is perturbed, retention factors and poly(A)+ RNA preferentially segregate with SON-rich structures. This association is notable in light of evolutionary analyses showing that SON IDRs expanded with the emergence of nuclear speckles in amniotes and are essential for formation of functional speckles ^13^. Recent work further identified THRAP3/BCLAF1-containing perispeckle networks that extend into the interchromatin space and associate with active genes ^30^, potentially providing a route for mRNP transfer from transcription sites to nuclear speckles.

The sponge-like architecture we observe may provide an efficient solution to mRNP retention. An extended scaffold–lumen interface could provide abundant docking sites for retention factors while leaving luminal space available to accommodate the mRNPs. These RNA-rich luminal spaces expand and contract with RNA load, allowing the architecture to accommodate changing amounts of mRNPs. Liao and Regev proposed that preferential partitioning of exons into nuclear speckles and introns toward the nucleoplasm positions splice sites at the speckle interface and thereby facilitate splicing ^40^. Although proposed for the outer speckle boundary, the same principle could operate throughout the internal scaffold network, greatly expanding the available RNA-scaffold interface for retention-mediated pre-mRNP quality control.

## Supporting information

smFISHprobes

## Resource availability

### Lead contact

Further information and requests for resources and reagents should be directed to the corresponding author, Maria Hondele, and will be fulfilled upon reasonable request.

### Material availability

Plasmids, oligonucleotides and cell lines are available through the lead contact upon request.

### Data and Code availability

- Imaging data will be deposited in the BioImage Archive (EMBL-EBI).
- All mass spectrometry proteomics data associated with this manuscript have been deposited to the ProteomicsXchange consortium via MassIVE (https://massive.ucsd.edu) with the accession numbers MassIVE MSV000103134/ PRIDE PXD083777. (Dataset is currently private. Reviewers can access the dataset following the instructions on the website. Username: MSV000103134_reviewer, Password: PCF).
- Any additional information required to reanalyze the data reported in this paper is available from the lead contact upon request.

## Supplemental Information

Supplemental Table 1: smFISH probes used in this study

## Acknowledgements

We are grateful to Claudia Keller Valsecchi, Tobias D. Williams and all members of the Hondele lab, particularly Daan Overwijn, Michelle Gut, and Ferdinand Weidner, for stimulating discussions and critical feedback on the manuscript. We further thank Jonas Bürki, Victoria Hilgers, Mirjam Uhland, and Tamara Utzinger for their valuable assistance in experimental work during their study internships and apprenticeships. We further thank the Imaging Core Facility (IMCF) and Proteomics Core Facility (PCF) of the Biozentrum, University of Basel, for their expert support throughout this project.

M.H. was supported by the Swiss National Science Foundation (PCEFP3_187052), the European Research Council (ERC-ST2020 950262), and institutional funds allocated to the Biozentrum, University of Basel, by the cantons of Basel-Stadt and Basel-Landschaft. K.D. was supported by the University of Basel’s Research Fund for Junior Researchers. F.V. acknowledges funding by the Swiss National Science Foundation PRIMA grant (PR00P3_208595). Funding for J.U. was supported by the Wellcome Trust (340926/Z/25/Z) and the UK Dementia Research Institute (UK DRI-RE21605) through UK DRI Ltd, which is principally funded by the UK Medical Research Council.

## Author contributions

conceptualization, K.D., and M.H.;

formal analysis, K.D., K.K., N.Beer, M.K.J;

investigation, K.D., K.K., N.Beu., N.Beer, K.D., L.S., D.H., G.B., S.L.;

methodology, K.D., A.F and M.K.J;

resources, K.D., K.K., N.Beu., L.S. and D.H;

writing - original draft, K.D. and M.H.;

writing - reviewing & editing, J.U. and F.V.;

visualization, K.D., N.Beer, M.K.J. and M.H.;

supervision, K.D., J.U., F.V. and M.H;

funding acquisition, K.D., J.U., F.V. and M.H.

## Declaration of generative AI and AI-assisted technologies in the writing process

During the preparation of this work, the author(s) used ChatGPT and CLAUDE.ai to improve wording and the clarity of the manuscript and to assist with script writing. The author(s) reviewed and edited the content as appropriate and take full responsibility for the content of the publication.

## Supplemental information

Supplemental Table 1: smFISH probes used in this study

## Material and Methods

### Key resources table

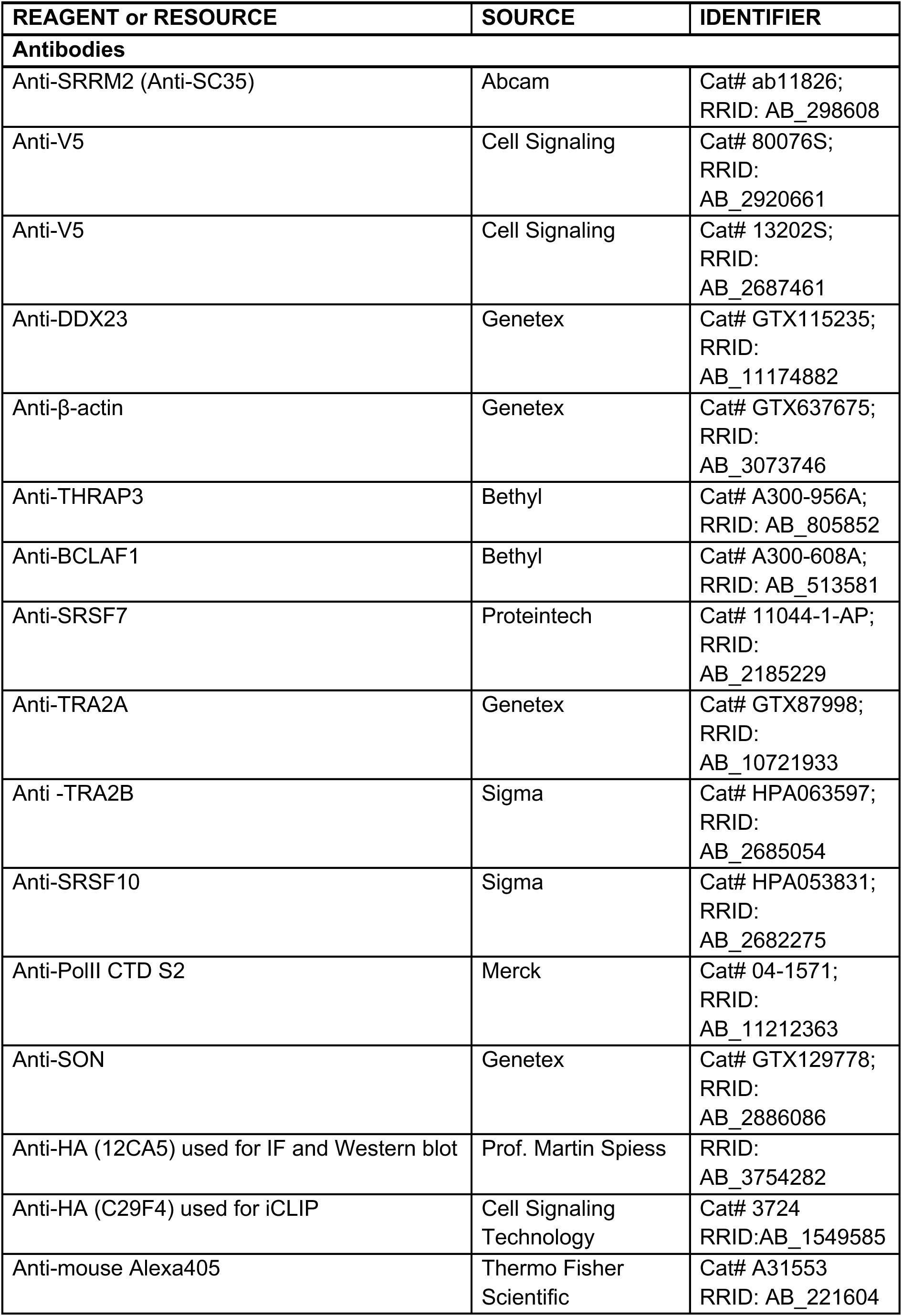
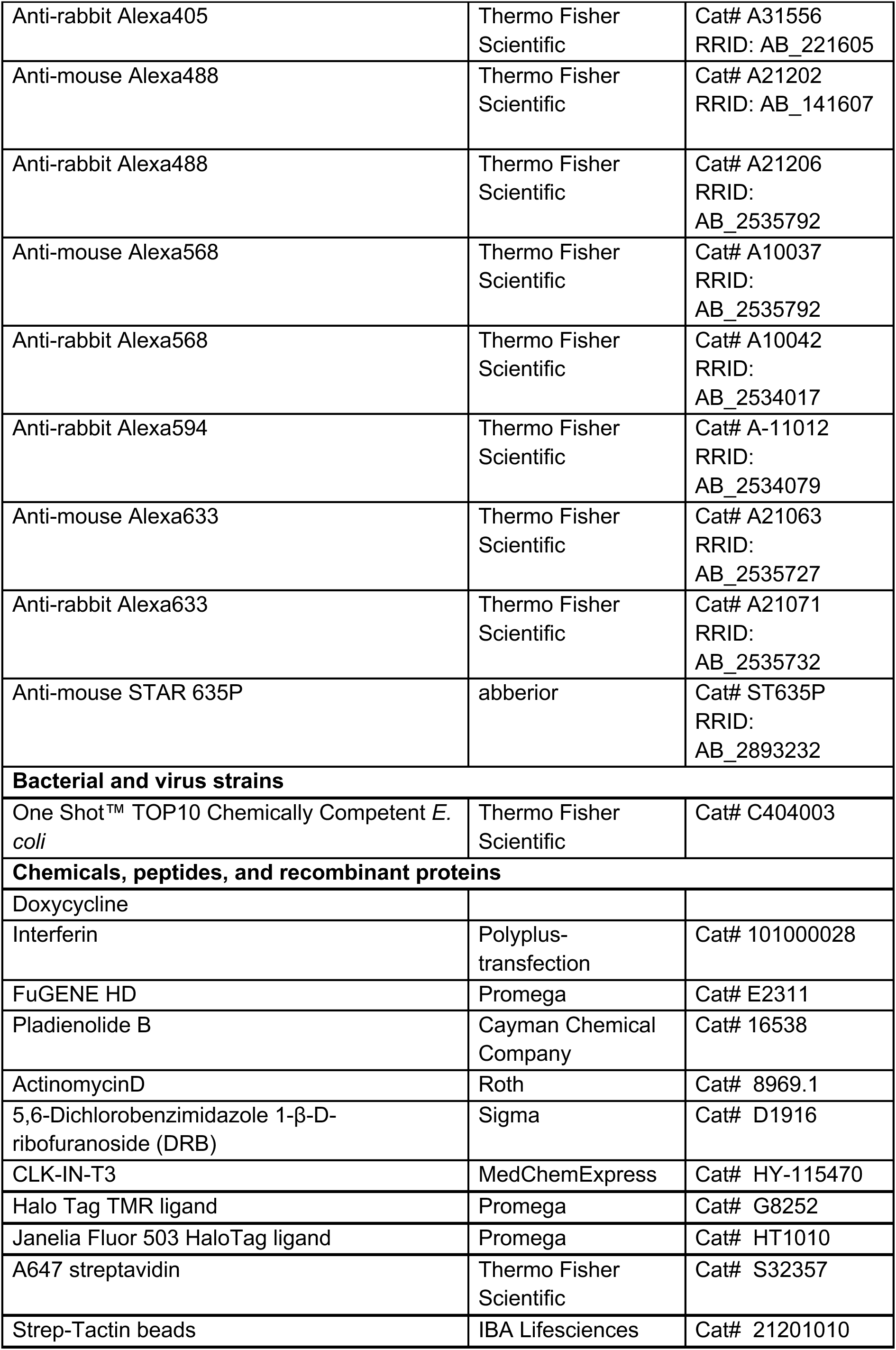
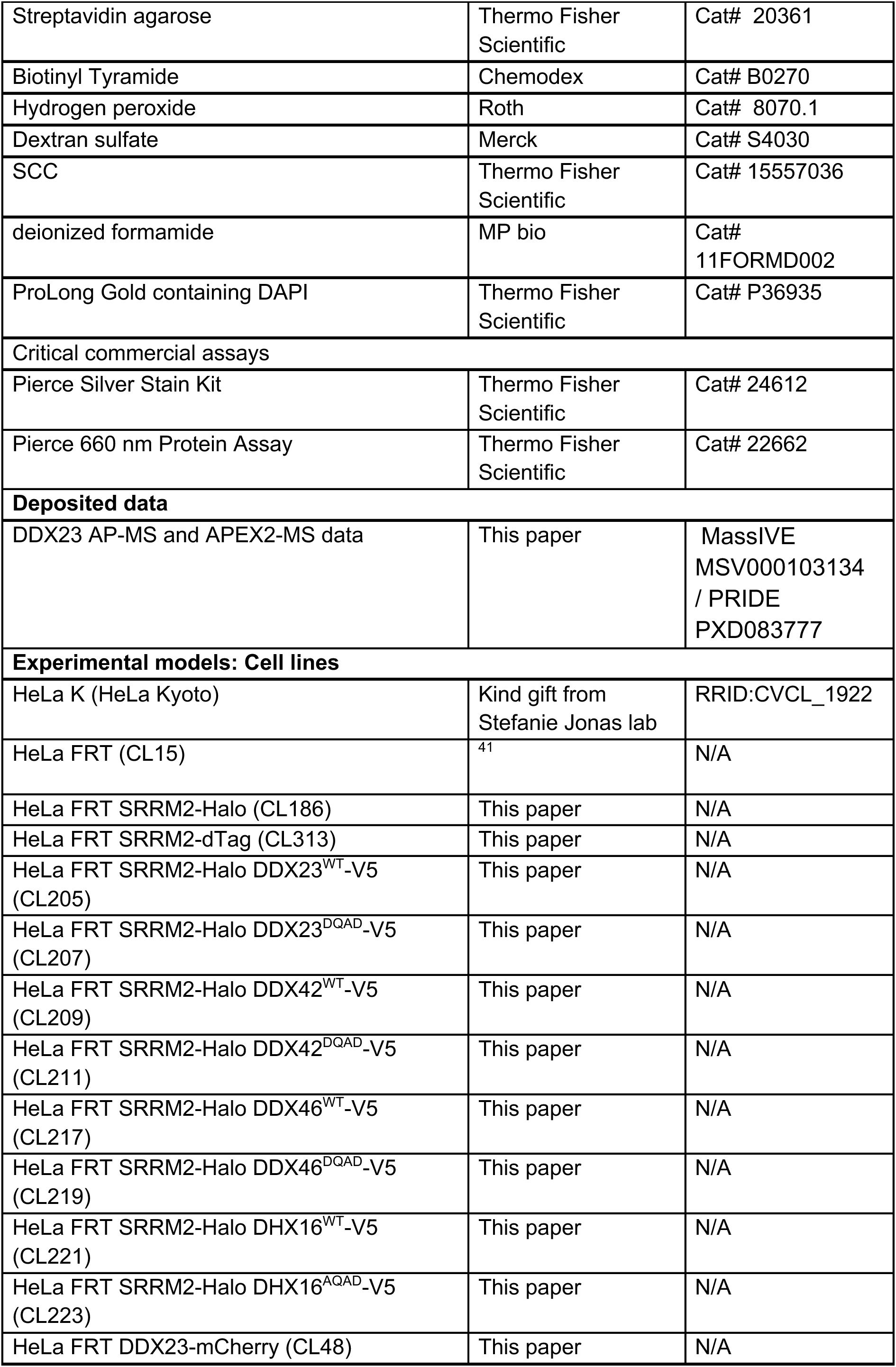
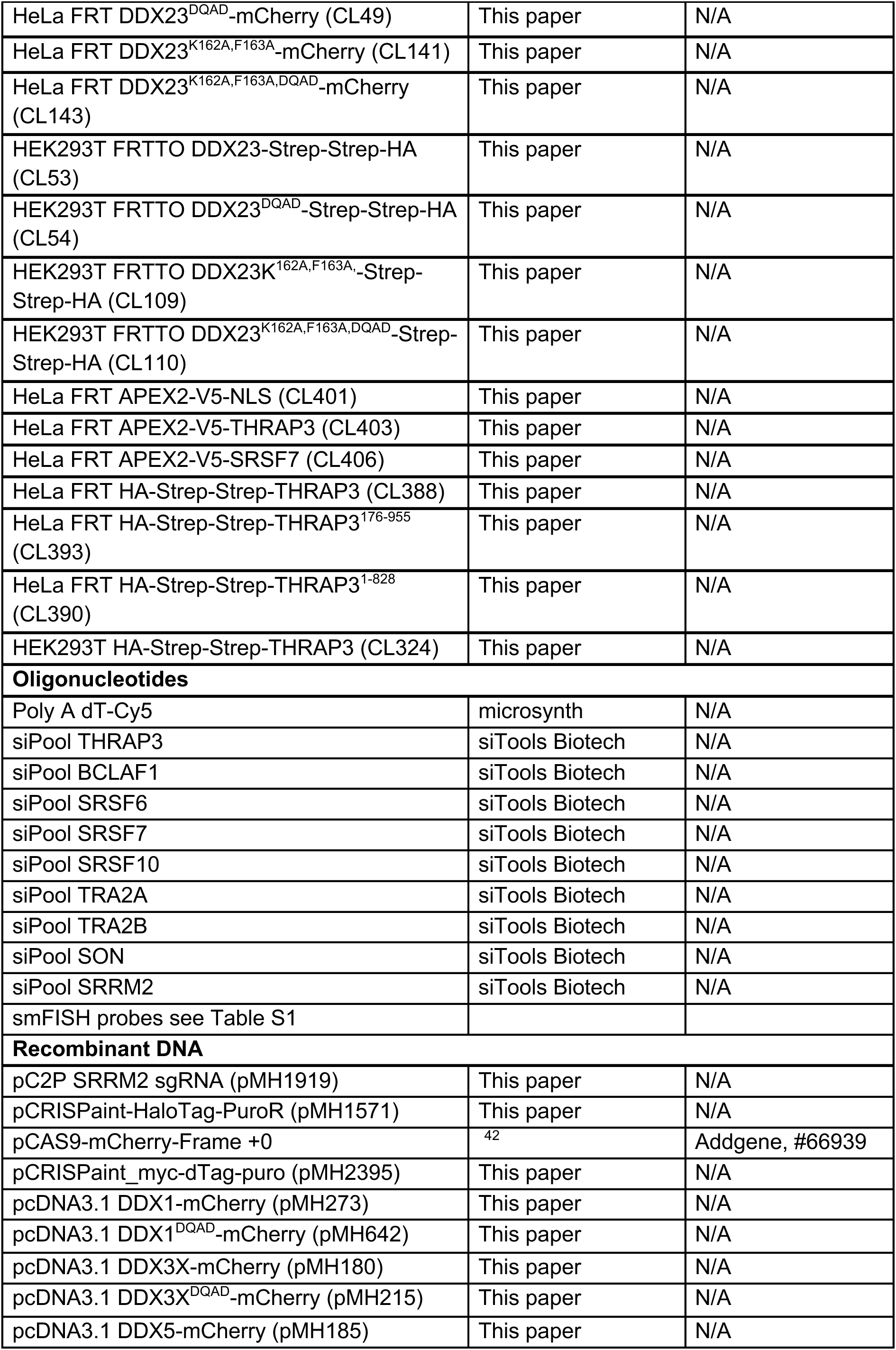
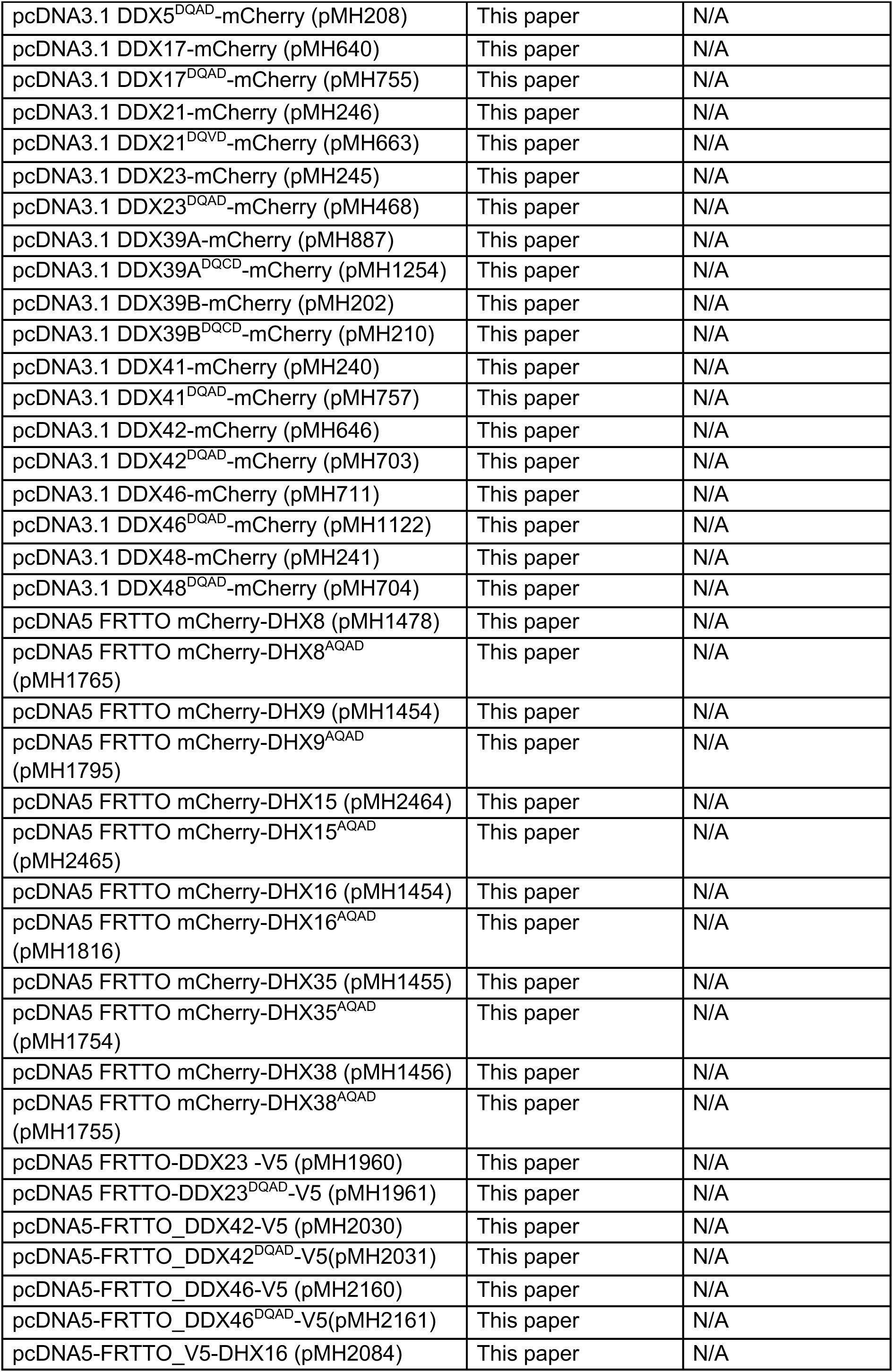
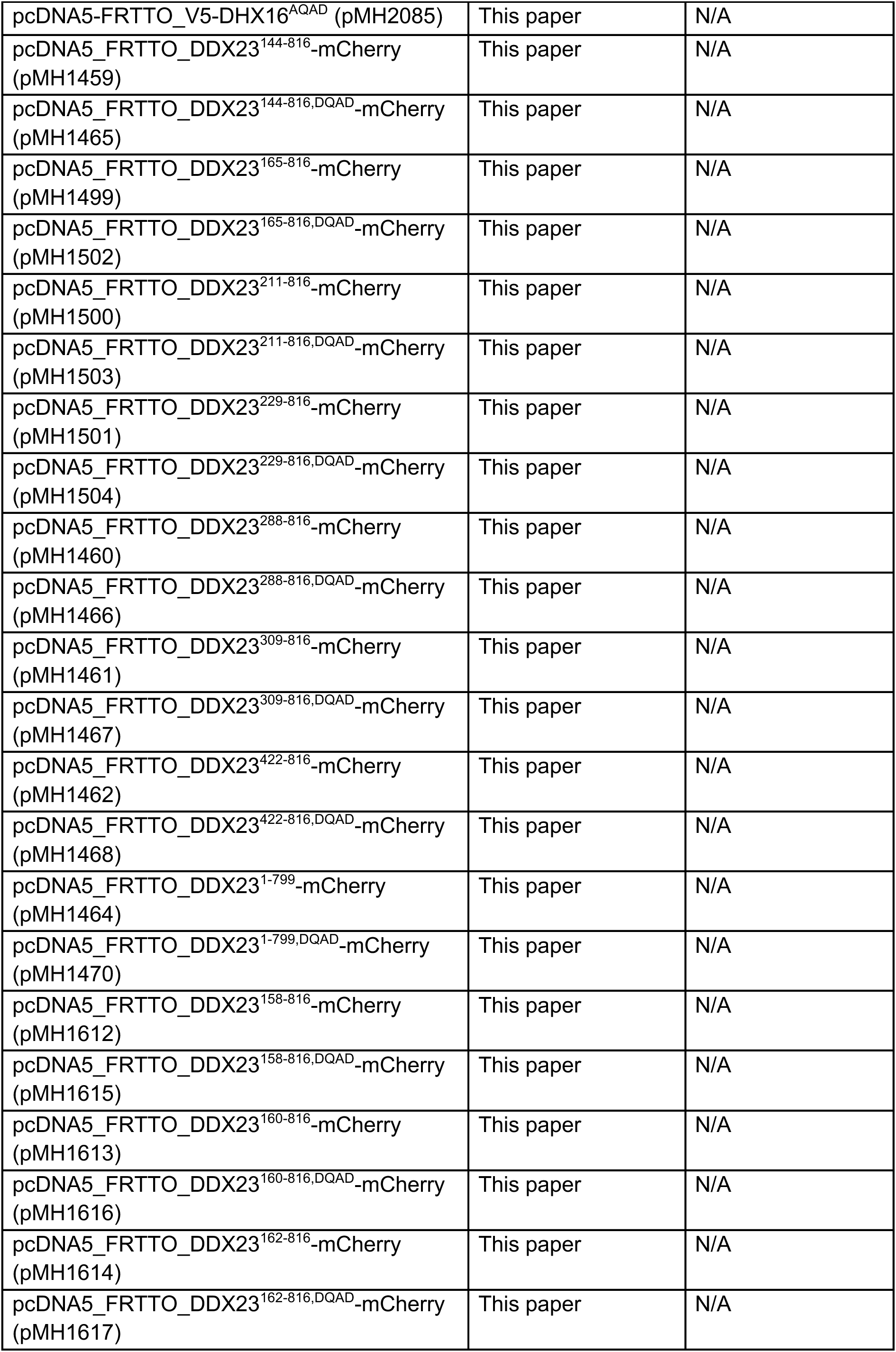
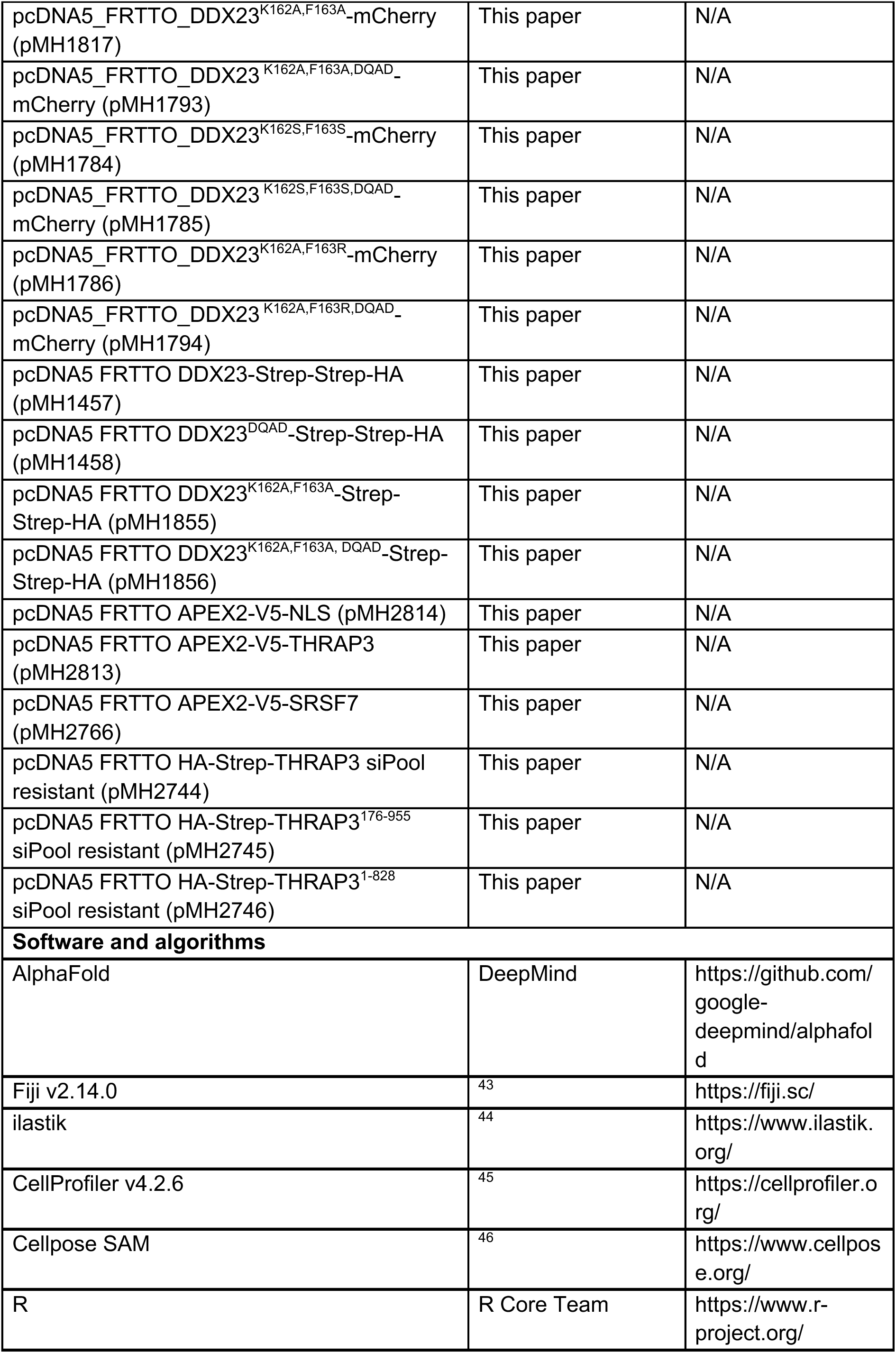
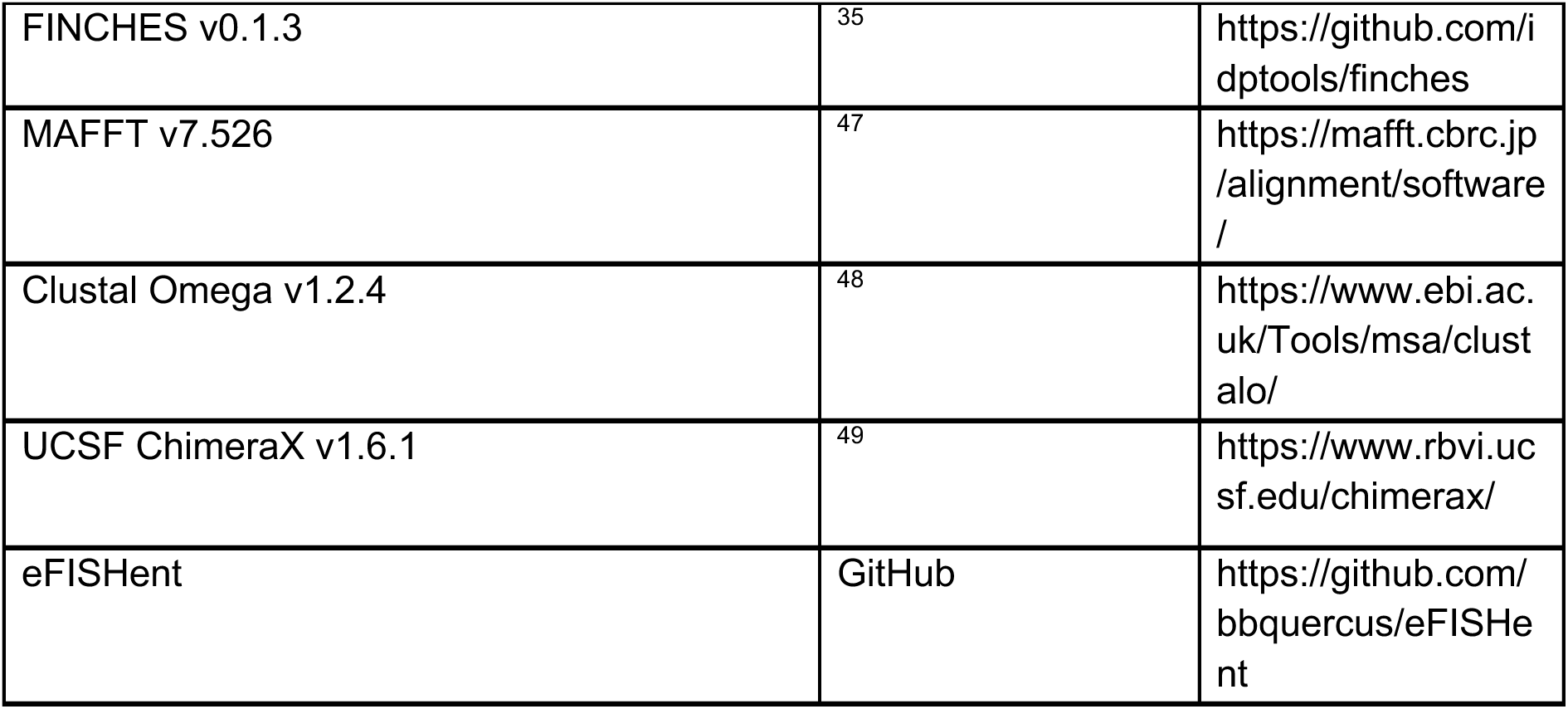

### Cell line maintenance and generation

Cell lines were maintained at 37°C with 5% CO_2_ in DMEM supplemented with 10% fetal bovine serum and 100 μg/mL penicillin/streptomycin. Cells were passaged before reaching confluence and were routinely tested for mycoplasma contamination. Plasmid transfections were performed using FuGENE HD according to the manufacturer’s instructions.

Endogenously tagged SRRM2 cell lines were generated in a parental HeLa FRT background ^41^ using the CRISPaint homology-independent knock-in strategy ^42^. A +0 Cas9/frame-selector plasmid, a guide-RNA plasmid targeting the SRRM2 stop codon (guide sequence: CCATGAGACACCGCTCCTCC) and a donor plasmid were co-transfected. The donor plasmid encoded HaloTag followed by a T2A-linked puromycin-resistance cassette. 48 h after transfection, cells were selected with 0.6 µg/mL puromycin until resistant colonies were obtained, from which single-cell-derived clones were isolated and expanded. Correct integration was confirmed by insertion-junction PCR followed by Sanger sequencing. Homozygous knock-in clones were identified using a separate PCR assay spanning the unmodified SRRM2 locus to test for the absence of the wild-type allele. SRRM2-Halo was labelled by incubation in medium containing 100 nM TMR-Halo dye for 18 h or 2.5 h before fixation.

Inducible cell lines were generated using the Flp-In system (Invitrogen) through stable integration of pcDNA5-based constructs. HeLa FRT cells, HeLa FRT SRRM2–Halo cells, or HEK293T FRT cells carrying a genomic FRT site were used as parental cell lines. Starting 48 h after transfection, cells were selected with hygromycin B at 0.3 mg/mL for HeLa cells or 0.1 mg/mL for HEK293T cells. Monoclonal cell lines were generated from HeLa cells, whereas polyclonal cell populations were established for HEK293T cells. Transgene expression was induced by the addition of doxycycline. Where indicated cells were treated with the following drugs: pladienolide B (500 nM, 2.5 h), actinomycin D (1 μg/mL, 2.5 h), DRB (50 µg/mL, 2.5 h), CLK inhibitor CLK-IN-T3 1 µM, 8 h or 24 h).

### RNA interference

Proteins of interest were depleted using siRNA pools purchased from siTOOLs Biotech, each containing 30 individual siRNAs to minimize off-target effects. Cells were seeded in 6-well plates at densities chosen such that siCTRL-treated cells reached approximately 80–90% confluence at the end of the respective depletion period. siRNAs were transfected in 200 µL Opti-MEM containing the indicated amount of siRNA and 2 µL INTERFERin, according to the manufacturer’s instructions.

The following final siRNA concentrations and depletion durations were used: siSRSF7, 3 nM for 72 h; siSRSF6, 3 nM for 72 h; siSRSF10, 3 nM for 72 h; siBCLAF1, 3 nM for 48 h; siTHRAP3, 3 nM for 48 h; siTRA2A, 1 nM for 48 h; siTRA2B, 1 nM for 48 h; siSON, 3 nM for 48 h; and siSRRM2, 15 nM for 72 h. For combined knockdowns, each siRNA pool was used at the same final concentration as in the corresponding individual depletion condition.

### Immunoblotting

Cells were harvested in SDS sample buffer containing β-mercaptoethanol and heated at 95°C for 5 min and treated in an ultrasonic bath for 10 min. Proteins were separated on 8–17% gradient SDS–PAGE gels in running buffer containing 25 mM Tris, 192 mM glycine and 0.1% SDS. Proteins were transferred to PVDF membranes either by wet transfer using buffer containing 25 mM Tris and 192 mM glycine, or by semi-dry transfer using buffer containing 33.6 mM Tris, 26 mM glycine, 14 mM tricine and 0.25 mM EDTA. For SRRM2 immunoblotting, cells were resuspended in SDS sample buffer containing β-mercaptoethanol, heated at 95°C for 5 min and sonicated in an ultrasonic bath for 10 min. Proteins were separated on 3–8% NuPAGE Tris-Acetate gels (Invitrogen) in Tris-Acetate SDS Running buffer (invitrogen) at 150 V for 50 min at RT and then transferred to an activated PVDF membrane in 25 mM Tris and 192 mM glycine, at 60 V for 3.5 h at 4°C.

For chemiluminescent detection, membranes were blocked in 4% milk in PBST, incubated with primary antibodies overnight at 4°C, washed 3 times in PBST and subsequently incubated with HRP-conjugated secondary antibodies. After three washes with PBST signals were developed using ECL and detected on a Fusion imaging system (Vilber). For near-infrared fluorescent detection, PVDF membranes were air-dried following transfer and reactivated in methanol before blocking in Intercept TBS Blocking Buffer (LI-COR). Membranes were incubated with primary antibodies overnight at 4°C, washed three times in TBS-T and subsequently incubated with IRDye-conjugated secondary antibodies. After three washes with TBS-T, fluorescent signals were detected using an Odyssey Fc Imaging System (LI-COR).

### Immunofluorescence analysis

Cells grown on coverslips were fixed with 3 or 4% paraformaldehyde in PBS for 15 min at room temperature and washed three times with PBS. Cells were permeabilized with 0.1% Triton X-100 and 0.02% SDS in PBS for 7 min, followed by blocking with 2% BSA in PBS for 30 min at room temperature. Primary antibodies were diluted in blocking solution and incubated with the cells for 1-2 h at room temperature or overnight at 4°C. Cells were then washed three times for 5 min with PBS and incubated with the appropriate secondary antibodies, diluted in blocking solution, for 1 h at room temperature. After three washes with PBS, cells were stained with DAPI where applicable, washed once more with PBS and mounted onto glass slides using Vectashield mounting medium and sealed with nail polish.

For SIM imaging, secondary antibodies conjugated to Alexa Fluor 405 or Alexa Fluor 488 were used, diluted 1:300; for STED imaging, secondary antibodies conjugated to STAR 635P or Alexa Fluor 594 were used diluted 1:150.

### Poly(A) RNA fluorescence in situ hybridization

Cells grown on coverslips were washed with PBS, fixed with 4% paraformaldehyde in PBS for 10 min at room temperature and washed four times with PBS. Cells were permeabilized in 70% ethanol for 2.5 h or overnight and washed with PBS containing 10% formamide for 5 min. Coverslips were incubated overnight at 37°C in hybridization buffer containing 0.5% BSA, 10% dextran sulfate, 2× SSC, 10% formamide, 200 nM yeast tRNA and 1 ng/µL Cy5-labelled poly(dT) probe. Cells were subsequently washed three times for 30 min with 2× SSC containing 10% formamide, stained with DAPI in 2x SSC, where applicable, mounted onto glass slides using Vectashield mounting medium and sealed with nail polish.

### Poly(A) RNA FISH combined with immunofluorescence

Cells grown on coverslips were washed with PBS, fixed with 4% paraformaldehyde in PBS for 10 min at room temperature and washed three times with PBS. Cells were permeabilized on ice for 10 min in PBS containing 0.5% Triton X-100 and 2 mM vanadyl ribonucleoside complex. After washing twice with PBS and twice with 2× SSC, cells were incubated overnight at 37°C in hybridization buffer containing 2× SSC, 10% formamide, 10% dextran sulfate, 10 mM DTT and 5 nM Cy5-labelled dT30 probe or Atto488 labelled dT30 probe. Cells were washed for 30 min at 37°C with 2× SSC containing 10% formamide, followed by washing three times with PBS and blocking with 1% BSA in PBS for 30 min. Cells were then incubated with primary antibodies for 1–2 h and secondary antibodies for 30 min at room temperature. Where applicable, nuclei were stained with DAPI for 10 min. Coverslips were washed with PBS and mounted onto glass slides using Vectashield and sealed with nail polish.

### Microscopy of immunofluorescence and Poly(A) RNA FISH

#### Confocal microscopy

Confocal images were acquired using a Zeiss LSM700 point-scanning confocal microscope equipped with a 63×/1.4 NA Plan-Apochromat oil-immersion objective or a Nikon Ti2 CrEST X-Light V3 spinning-disk confocal microscope equipped with either a 60×/1.4 NA or a 100×/1.45 NA CFI Plan Apochromat Lambda D oil-immersion objective.

#### Structured illumination microscopy (3D-SIM)

3D-SIM imaging was performed as follows. Fixed cells prepared for super resolution microscopy were imaged using a Deltavision OMX-Blaze V4 microscope system (Applied Precision/Leica)equipped with a 60×/NA 1.42 Plan Apo N oil immersion objective, four liquid-cooled sCMOScameras, and laser lines at 405, 488, 568, and 642 nm. Images were acquired as z-stacks with 2 to 4.5 μm total z-range and 0.125 μm z-step size. Excitation light was directed through a movable optical grating to generate a fine-striped interference pattern on the sample plane. The pattern was shifted laterally through five phases and three rotations of 60° for each z section. The exposure time was set to reach at least 2000 count.Raw images were reconstructed and channels were aligned using the Deltavision OMX softWoRx softwarepackage (Applied Precision).

#### TauSTED microscopy

STED imaging was performed on a Leica STELLARIS 8 FALCON microscope using an HC PL APO 100×/1.40 NA oil-immersion STED WHITE objective. Fluorescence was detected with a HyD X detector, and 2D STED was achieved using a 775-nm depletion laser. Image acquisition was performed in LAS X, using the FALCON lifetime modality with 10 line accumulations, and TauSTED/Xtend processing were applied post-acquisition. Images were processed with the following parameters (background suppression/tau-strength/denoise/Xtend strength/Xtend smooth/Xtend low intensity) were used: Figure S12C: SRRM2 (yes/50/70/5/0/20), SON (yes/100/80/5/0/20). Figure S12H: SRRM2 (yes/40/60/5/0/40), SON (yes/50/60/5/0/0).

#### Airyscan microscopy

Airyscan imaging was performed on an inverted Zeiss LSM880 point-scanning confocal microscope equipped with an AiryScan 32-subunit detector and a 63×/1.40 NA Plan-Apochromat oil-immersion objective. Images were processed using the Airyscan super-resolution reconstruction algorithm in ZEN Blue software, strength settings SRRM2 6.2, PolyA 6.4, SON 6.8.

#### Image processing and quantitative image analysis

Image analysis of confocal images was performed using Fiji, ilastik and CellProfiler. For z-stack acquisitions, maximum-intensity projections were generated in Fiji using a custom macro prior to analysis. Nuclear speckles were identified using the supervised Pixel Classification workflow in ilastik, and the resulting probability maps were imported into CellProfiler for segmentation. Nuclei were segmented in CellProfiler and background correction was performed prior to quantitative intensity measurements. Where required for poly(A) RNA analysis, cytoplasmic regions were additionally defined in CellProfiler.

Quantitative measurements, including nuclear speckle number, area and fluorescence intensity, as well as nuclear and nuclear speckle fluorescence intensities of proteins and poly(A) RNA, were extracted on a per-cell basis for downstream analysis. For poly(A) RNA, cytoplasmic fluorescence intensity was additionally quantified where required.

### 3D-SIM and TauSTED image processing and quantitative image analysis

Reconstructed 3D-SIM volumes were analysed volumetrically in Python (NumPy, SciPy, scikit-image, statsmodels); no projections were used. Reconstruction negatives were clipped to zero, each channel was denoised with a Gaussian of σ = 40 nm in physical units and scaled by its own 99.9th percentile, so that all reported quantities are ratios or fractions. Nuclei were segmented by Otsu thresholding of the scaffold signal smoothed with σ = 1 µm. Speckle bodies were defined by Otsu thresholding at σ = 200 nm within the nucleus, followed by closing and hole filling, intersected with an 80 nm dilation of the scaffold-node mask obtained by a second Otsu threshold at σ = 42.5 nm (100 nm FWHM); objects below 0.05 µm³ were discarded. Within bodies, scaffold voxels were defined by per-channel Otsu thresholds and the remainder termed interstitium; the core comprises body voxels more than 80 nm inside the body surface, and the axially trimmed core additionally excludes two z-planes from each end of every object.

Overlap between the two scaffold markers, and between poly(A) RNA and the scaffold, was quantified after matching occupancy, thresholding each channel at its own top 40 % of body voxels, fixing the chance Jaccard index at 0.25; a per-cell null was obtained by rotating one channel 90° in xy, giving 0.249–0.258. Lateral cross-correlations were computed by FFT within the body mask, corrected for mask shape and normalised so that zero lag equals the Pearson coefficient. Interstitial gap calibre and strand thickness lie at the sampling limit in all conditions and are not reported.

RNA distribution was measured as mean poly(A) intensity in body voxels binned by distance from the scaffold surface (0–40, 40–80, 80–120, 120–200 nm; normalised to the scaffold of the same cell. Interstitial occupancy was evaluated across thresholds set at 4 times each cell’s nucleoplasmic median.

### Single-molecule fluorescence in situ hybridization (smFISH)

#### Probe design

smFISH probe sets were designed as pools of short single-stranded DNA (ssDNA) oligonucleotides tiling the target transcript, each subsequently labelled at its 3′ end with a single fluorophore (see *Probe labelling*). 50 nt probes against BRD4 targeted exonic (mature mRNA) sequences and were as described in ^17^. 40-50 nt probes against KPNA1 were designed against both exonic and intronic sequence using eFISHent (https://github.com/bbquercus/eFISHent/wiki) using hg38 as reference.

#### Fluorophore–ddUTP conjugation

For conjugation of amino-11-ddUTP (Lumiprobe, #15040) to NHS-ester dyes (Atto 565-NHS, ATTO-tec AD 565-31; and/or Atto 633-NHS, ATTO-tec AD633-31), all steps were performed in a moisture-free chamber (regenerated silica gel, ≥1 h equilibration). Dye-NHS esters were reconstituted to 40 mM in anhydrous DMSO (Atto 565; Atto 633). Per reaction, 5 mM amino-11-ddUTP, 10 mMdye-NHS ester and 50 mM NaHCO₃ pH 8.4 were mixed and incubated for 2 h at room temperature protected from light. The reaction was then diluted with nuclease-free water, yielding dye-conjugated ddUTP at 5 mM. Conjugated nucleotides were stored at −20 °C.

#### Probe labelling and purification

Probe pools were labelled at the 3′ end by terminal deoxynucleotidyl transferase (TdT)-mediated addition of a single dye-conjugated ddUTP, following a protocol adapted from ^50^. Oligonucleotides were first combined into a 100 µM equimolar probe mix. Per labelling reaction: 63 µM probe mix, 1× TdT buffer (200 mM potassium cacodylate, 25 mM Tris, 0.01% (v/v) Triton X-100, 1 mM CoCl2 (pH 7.2)), 0.33 mM dye-ddUTP and 1.3 units/µL TdT (Thermo Fisher, EP0162) were combined and incubated at 37 °C for 24 h in a thermocycler.

Labelled probes were purified by ethanol precipitation on magnetic beads using a KingFisher system. To each reaction, 60 µL of 1 M Na-acetate (pH 5.5), 115 µL nuclease-free water, and 1.5 µg linear acrylamide (Thermo Fisher, AM9520) were added. The resulting mixture was transferred to a KingFisher 96-well plate (Thermo Fisher, 95040450) containing 800 µL ethanol supplemented with 10 µL carboxylated magnetic beads (Thermo Fisher, 65011). Following precipitation for 1 h at room temperature under constant mixing, the probes were washed twice with 80% ethanol and eluted in 30 µL nuclease-free water. Eluates were kept nuclease-free and protected from light.

Labelling efficiency was assessed by NanoDrop absorbance spectroscopy (A₂₆₀ for nucleic acid; A₅₇₀ for Atto 565 and A₆₃₄ for Atto 633; values corrected for the 1 mm NanoDrop pathlength). Oligo and dye concentrations and the degree of labelling (DOL = c_dye / c_oligo) were calculated from the dye-corrected A₂₆₀ and the dye absorbance using the manufacturer-provided molar extinction coefficients, with the oligo extinction coefficient increased by 9,000 M⁻¹ cm⁻¹ to account for the added 3′ terminator nucleotide. Purified, labelled probes were stored at −20 °C protected from light.

#### Cell culture and fixation

For smFISH, 12 mm #1.5 precision glass coverslips were placed in 6-well plates and seeded with 6 × 10⁴ cells per well, then grown for 3 days. Cells were washed 1–2× with 1× nuclease-free PBS and fixed in 4% paraformaldehyde in 1× PBS for 10 min at room temperature with gentle agitation. Fixed cells were washed twice with PBS and permeabilised in 70% ethanol at 4 °C overnight.

#### Hybridisation, washing and mounting

Hybridisation buffer was prepared fresh immediately before use and contained 10% (w/v) dextran sulfate, 2× SSC and 45% (v/v) formamide, with each probe added to a final concentration of 125 nM. Wash buffer contained 2× SSC and 45% (v/v) formamide. Formamide was equilibrated to room temperature before use. Following permeabilisation, coverslips were washed twice for 5 min in wash buffer. Each coverslip was then inverted (cells facing down) onto a 40 µL droplet of probe-containing hybridisation buffer in a humidified chamber, sealed, and incubated in the dark at 37 °C for 4 h to overnight. Coverslips were then washed three times for 30 min at 37 °C in pre-warmed wash buffer with gentle agitation, followed by a single PBS wash. Coverslips were mounted onto glass slides with ProLong Gold containing DAPI and sealed with nail polish. Images were acquired on a Visitron VisiScope spinning-disk confocal system consisting of a Nikon Eclipse Ti-E inverted microscope equipped with a Yokogawa CSU-W1 scan head and an EMCCD camera. Excitation was provided by 405, 488, 561, and 640 nm solid-state lasers. Images were acquired using either a 60×/1.4 NA or 100×/1.4 NA oil-immersion objective.

#### ImageProcessing

Nuclear segmentation was performed using the Cellpose-SAM cpsam model on the DAPI channel. Speckles were segmented by automatic Otsu-based thresholding. Cell segmentation was performed using the Cellpose-SAM cpsam model on smFISH images after Gaussian filtering (σ = 8) and rolling-ball background subtraction (radius = 500 pixels). Code for used for processing smFISH images can be found here: https://github.com/Voigt-Lab/NB_smfish-speckle-analysis

### Affinity purification followed by mass spectrometry

Stable HEK293T FRT/TO cell lines expressing HA–Strep–Strep-tagged DDX23 variants or HA–Strep–Strep–GFP as a control were induced with doxycycline for 8 h. Si× 10-cm dishes were used per cell line and biological replicate. Cells were harvested in PBS, washed once and lysed for 10 min at 4°C in 50 mM Tris-HCl, pH 7.5, 300 mM NaCl, 5 mM MgCl₂, 0.5% NP-

40 and 1 mM DTT supplemented with protease inhibitors. Lysates were clarified by centrifugation at 20,800 × g for 5 min at 4°C and incubated with pre-equilibrated Strep-Tactin beads for 30 min at 4°C with rotation. Beads were collected by centrifugation at 300 × g for 1 min and washed six times with 50 mM Tris-HCl, pH 7.5, 300 mM NaCl, 5 mM MgCl₂ and 1 mM DTT. Bound proteins were eluted for 4 min on ice using the same buffer supplemented with 2 mM biotin. Eluates were adjusted to final concentrations of 5% SDS, 100 mM triethylammonium bicarbonate (TEAB) and 10 mM TCEP, supplemented with 15 mM iodoacetamide, and incubated for 30 min at 25°C in the dark. Subsequently, protein digestion and desalting was performed by SP3 on a Tecan Fluent platform (Tecan) following the published procedure ^51^. Vacuum dried peptides were stored at -20 C before mass spectrometry analysis.

Mass spectrometry analysis was performed on a Orbitrap Exploris 480 platform as described elsewhere ^52^. Raw data was searched by MSFragger using FragPipe (version 22.0) ^53^ against the UniProt human protein database (version 22^nd^ February 2022) and common protein contaminants. Differential protein abundance analysis was performed using MSstats (version 4.13) ^54^.

#### Silver staining

Protein samples were separated by SDS–PAGE and visualized using the Pierce Silver Stain Kit according to the manufacturer’s instructions.

#### APEX2 proximity labeling and streptavidin enrichment followed by mass spectrometry

Stable cell lines expressing APEX2–V5 fused to SRSF7, THRAP3 or an NLS control were induced with doxycycline for 72 h. Four 10-cm dishes per condition were used for APEX2– SRSF7 and the APEX2–NLS control processed under the corresponding conditions, whereas four 15-cm dishes per condition were used for APEX2–THRAP3 and the respective APEX2– NLS control samples. Where indicated, cells were treated with 500 nM pladienolide B for 2.5 h before harvesting.

For proximity labeling, cells were incubated for 30 min in medium containing biotinyl tyramide. A final concentration of 0.5 mM biotinyl tyramide was used for APEX2–SRSF7 and the corresponding NLS control, whereas 1 mM was used for APEX2–THRAP3 and their corresponding NLS controls. When Pladienolide B was used it was maintained throughout the labeling procedure. Labeling was initiated by addition of 0.5 mM H₂O₂ for 1 min and quenched by three washes with PBS containing 10 mM sodium ascorbate, 10 mM sodium azide and 5 mM Trolox. Cells were collected by centrifugation at 3,000 × g for 10 min at 4°C, flash-frozen and stored at −80°C.

Cell pellets were lysed in RIPA (50 mM Tris-HCl pH 7.5, 150 mM NaCl, 0.1% SDS, 0.5% sodium deoxycholate, 1% Triton X-100) buffer supplemented with protease inhibitors, 1 mM PMSF, 5 mM Trolox, 10 mM sodium azide and 10 mM sodium ascorbate. Lysates were incubated on ice for 30 min, sonicated (15 pulses of 1s, 30s break, 15 pulses of 1s) and clarified by centrifugation at 15,000 × g for 15 min at 4°C. Protein concentrations were determined using the Pierce 660 nm Protein Assay, and lysates were equalized for protein content within each experimental set. Equal amounts of lysate were incubated with streptavidin agarose beads (Thermo Scientific, #20361) for 3 h at 4°C. Beads were washed twice with RIPA buffer, once with 1 M KCl, once with 0.1 M Na₂CO₃ and once with 2 M urea in 10 mM Tris-HCl, pH 8.0, and twice again with RIPA buffer. Biotinylated proteins were eluted by heating the beads for 10–15 min in 5% SDS, 10 mM TCEP, 100 mM triethylammonium bicarbonate and 2 mM biotin, flash-frozen and stored at −80°C until further processing for mass spectrometry.

Protein digestion and desalting was performed by SP3 as mentioned above. Dried peptide mixtures were dissolved in 0.1% formic acid (in water) and approximately 75 ng peptide mixture was analyzed on an Orbitrap Astral Zoom mass spectrometer which was equipped with a Vanquish Neo liquid chromatography system (both Thermo Scientific) and a custom-made column heater. A RP-HPLC column (75μm × 30cm) packed in-house with C18 material (ReproSil-Pur C18–AQ, 1.9 μm resin; Dr. Maisch GmbH) was used at a flow rate of 200 nL/min. The Orbitrap Astral Zoom was operated in DIA mode (100 windows, isolation window 6 m/z, range: 380-980 m/z, fill time 7 ms). DIA data was recorded in centroid mode. Loop control was set to 0.6 s. Raw data was searched by Spectronaut (version 19.0, Biognosys AG) against the UniProt human protein database (version 22^nd^ February 2022) and common protein contaminants. Differential protein data analysis was performed using the Proteoflux analysis pipeline ^55^.

### iCLIP

HEK293T cells were induced to express HA-THRAP3 at near endogenous level for 48h. Culture medium was removed and cells were washed with ice-cold PBS while maintained on ice. Cells were irradiated once with 150 mJ/cm² UV-C light at 254 nm using a Stratalinker 2400. Cells were harvested by scraping, transferred to tubes and pelleted at 500 × g for 3 min at 4°C. Cell pellets were snap-frozen and stored at −80°C until further processing. Four iCLIP replicates of HA-tagged THRAP were then performed according to the iCLIP protocol ^56^, using crosslinked cell lysate containing 1.5 mg of protein per sample. For immunoprecipitation, 5 μg of anti-HA (C29F4) antibody was used.

#### iCLIP and eCLIP analysis

iCLIP data were first demultiplexed on the Flow web server (https://app.flow.bio/) using the Demultiplex pipeline v1.1 (https://github.com/goodwright/flow-nf/tree/master/subworkflows/goodwright/demultiplex) run with Nextflow v24.04.2 ^57,58^. In this pipeline Ultraplex v1.2.9 (https://github.com/ulelab/ultraplex) is used with settings -m5 1 -m3 0 -q 30 -q5 0 -mt 3 -l 16 --adapter AGATCGGAAGAGCACACGTCTG to demultiplex the paired-end FASTQ files into individual sample FASTQ files, move the UMI to the read header and trim read 3’ ends for adapter and quality ^59^. Subsequently, the samples were analysed using CLIP-Seq v1.7 (https://github.com/goodwright/clipseq/tree/1.7) run with Nextflow v24.04.2 on Flow. In this pipeline, reads are first trimmed for any remaining 3’ end Illumina adapters and for quality using Trim Galore v0.6.7 with Cutadapt v3.4 (--paired -q 20 --length 10). Trimmed reads are pre-mapped using Bowtie v1.3.0 with settings -v 2 -m 100 --norc -- best --strata to an index containing the 45S rDNA cluster (NCBI Gene ID: 106631777), 5S rRNA (NCBI Gene ID 100169751) and all tRNA (http://gtrnadb.ucsc.edu/GtRNAdb2/genomes/eukaryota/Hsapi38/hg38-mature-tRNAs.fa).

Unmapped reads are then mapped to Human genome GRCh38 with Ensembl v109 annotation using STAR v2.7.9a with settings --readFilesCommand zcat --outSAMtype BAM Unsorted -- quantMode TranscriptomeSAM --outFilterMultimapNma× 100 --outFilterMultimapScoreRange 1 --outSAMattributes All --alignSJoverhangMin 8 --alignSJDBoverhangMin 1 –outFilterType BySJout --alignIntronMin 20 --alignIntronMa× 1000000 --outFilterScoreMin 10 -- alignEndsType Extend5pOfRead1 --twopassMode Basic. Uniquely mapped reads are further filtered from mapped BAM files using Samtools v1.17 (samtools view -q 5), before being deduplicated by UMICollapse v1.0.0 based on start position and UMI (Liu 2019). Crosslink sites were defined as the position immediately upstream of the read start using BEDTools v2.30.0. Peaks were called on individual samples using Clippy v1.5.0 with default settings.

The Clippy peak files of the four replicates were downloaded from the Flow project (https://app.flow.bio/projects/788995297969977723/) and used as input for the THRAP3 co-binding analysis with RBPeek.

Raw ENCODE eCLIP fastq files for each RBP analysed in both HepG2 and K562 were downloaded from www.encodeproject.org, and processed using the nf-core/clipseq (v1.0.0) pipeline to generate crosslink bed files ^58^.

#### THRAP3 binding analysis

HA-THRAP3 iCLIP peaks were called with Clippy v1.5.0. Overlapping peaks were merged across replicates on the same strand, and loci present in 2 or more replicates were retained as a single-nucleotide midpoint (25,814 loci). Mitochondrial and intergenic regions were removed and loci were split into exons (n = 18,918) or introns (n = 6,857) using a filtered Gencode v39 GTF. The inference loci were compared with 224 eCLIP samples from HepG2 and K562 cell lines using RBPeek (https://github.com/ulelab/RBPeek). For each panel sample peak, the cDNA count was distributed uniformly across its width, and signal from the same strand was measured across each position within ±100 nt of a THRAP3 locus. Signal was normalised by the panel samples total peak cDNA in exons or introns respectively (per million) to control for sequencing depth. The metaprofile of a dataset is the mean over all loci of log(1 + normalised signal) at each position. Datasets were ranked by the area under the metaprofile within ±10 nt of the THRAP3 inference loci, and the highest-ranked datasets are shown. Heatmaps show the strongest peak within ±10 nt of each locus for the top 10% of datasets (24 of 224).

### Ortholog retrieval and quality control

Human THRAP3 (Ensembl gene ENSG00000054118; canonical protein ENSP00000346634; UniProt Q9Y2W1; 955 aa) and BCLAF1 (ENSG00000029363; ENSP00000435210; UniProt Q9NYF8; 920 aa) were used as query references. Vertebrate orthologs were retrieved from Ensembl Compara via the REST API homology/id endpoint restricted to orthologues, so that orthology assignments were taken directly from Compara’s gene-tree reconciliation. For species with multiple orthologs, a single representative was retained per species (highest percent identity to the human query). Protein sequences were recovered by removing gap characters from the Compara alignment strings. Sequences shorter than 50% or longer than 150% of the human reference length were flagged as fragments or mis-annotations and excluded from alignment. After filtering, 188 THRAP3 and 132 BCLAF1 orthologs (132 species with both genes) were carried forward.

#### Multiple sequence alignment

Ortholog sets were aligned separately for each protein, and the combined THRAP3 + BCLAF1 set (320 sequences) was aligned together for the paralog comparison, using MAFFT v7.526 with the L-INS-i strategy (--localpair --maxiterate 1000). To confirm that the conservation profile was not an artifact of a single aligner, each set was independently realigned with Clustal Omega v1.2.4; and MAFFT L-INS-i alignments were used for all downstream analyses.

#### Per-residue conservation scoring

Column-wise conservation was computed as the Jensen–Shannon divergence between the observed amino-acid distribution and a BLOSUM62-derived background distribution, following ^60^, with a gap penalty that down-weights columns in proportion to their gap fraction. Alignment columns were mapped back to human reference residue positions, and scores were smoothed with a centered 15-residue sliding-window mean for display. Columns with a gap fraction above 0.5 were flagged as low-confidence.

#### Paralog-shared identity

From the combined THRAP3 + BCLAF1 alignment, each column occupied by both paralogs was scored as identical when the majority-cons^61^ensus residue of the THRAP3 orthologs matched that of the BCLAF1 orthologs. The resulting binary per-residue identity signal was mapped to human coordinates and smoothed with a centered 21-residue sliding-window mean to yield a continuous paralog-identity track; contiguous residue runs exceeding 0.5 were defined as paralog-shared blocks.

#### Prediction of THRAP3–partner interactions with FINCHES

Sequence-encoded intermolecular interactions were predicted using FINCHES (v0.1.3) ^35^, which estimates a mean-field interaction parameter (ε) between disordered protein regions from residue chemistry. Canonical human sequences for THRAP3 (UniProt Q9Y2W1, 955 aa), SON (P18583, 2426 aa), and SRRM2 (Q9UQ35, 2752 aa) were obtained from UniProt. Because SON and SRRM2 exceed the length scale of a single FINCHES calculation, each was divided into non-overlapping fragments of ∼250 residues, with fragment boundaries shifted to avoid the few predicted folded regions (contiguous stretches ≥ 20 residues with disorder score < 0.5); THRAP3 was analyzed as a single full-length sequence (the common query). For each THRAP3 × fragment pair, a residue-resolved interaction matrix was computed with the FINCHES Mpipi ^61^ frontend using intermolecular_idr_matrix with a sliding window of 31 residues, together with the scalar mean-field ε from epsilon. By convention ε < 0 denotes net attraction and ε > 0 net repulsion. Fragment matrices were concatenated along the partner axis to reconstruct full-length THRAP3 × SON and THRAP3 × SRRM2 interaction maps. Per-partner summary statistics (mean ε across fragments; attractive contact area, defined as the fraction of matrix cells with ε < 0) were used to compare the two partners, and per-THRAP3-residue interaction profiles were obtained as the mean ε across the full partner protein (row means of the reconstructed maps).

#### General statistical analysis

Statistical analyses and data visualization were performed in R. Unless otherwise indicated, experiments were performed in three independent biological replicates (N = 3), with individual cells analyzed within each replicate. Biological replicates, rather than individual cells, were considered the unit of replication for statistical inference. Exact numbers of biological replicates and cells analyzed are provided in the corresponding figure legends.

Single-cell measurements are displayed as individual points together with boxplots and biological-replicate summaries. Boxplots show the median (center line) and interquartile range (IQR; box), with whiskers extending to the most extreme values within 1.5× IQR. Where shown, larger overlaid points represent the mean of each biological replicate. Unless otherwise indicated, comparisons between matched experimental conditions were performed using two-tailed paired t-tests on biological-replicate means unless otherwise indicated. Statistical tests specific to individual analyses are indicated in the corresponding figure legends. Statistical significance was defined as *p* < 0.05; ns, not significant.

### Image-based quantification

Nuclear speckle morphology was quantified on a per-cell basis. Nuclear speckle area was calculated as the mean area of segmented nuclear speckles within each nucleus, and nuclear speckle number as the number of segmented speckles per nucleus. Protein enrichment in nuclear speckles was quantified as the ratio of fluorescence intensity within nuclear speckles to the corresponding nuclear or nucleoplasmic fluorescence intensity, as indicated for the individual experiment. Where indicated, enrichment ratios were normalized within each biological replicate to the corresponding control condition and log₂-transformed.

### RNA localization analysis

Poly(A) RNA localization was quantified from background-corrected fluorescence intensities within the indicated cellular compartments. Depending on the experiment, poly(A) RNA accumulation was expressed as nuclear speckle enrichment relative to the surrounding nucleoplasm or as the nuclear-to-cytoplasmic fluorescence-intensity ratio. For selected experiments containing multiple levels of sampling, statistical significance was assessed using hierarchical linear mixed-effects models accounting for biological replicate and imaging field, followed by planned contrasts with correction for multiple comparisons as indicated in the corresponding figure legends.

For smFISH experiments, RNA enrichment in nuclear speckles was calculated from background-corrected speckle-to-nucleoplasm fluorescence-intensity ratios on a per-cell basis. Values were normalized within each biological replicate to the corresponding control condition and log₂-transformed where indicated. Statistical comparisons were performed on biological-replicate means using two-tailed paired t-tests.

## Supplementary figures and figure legends

**Figure S1.**
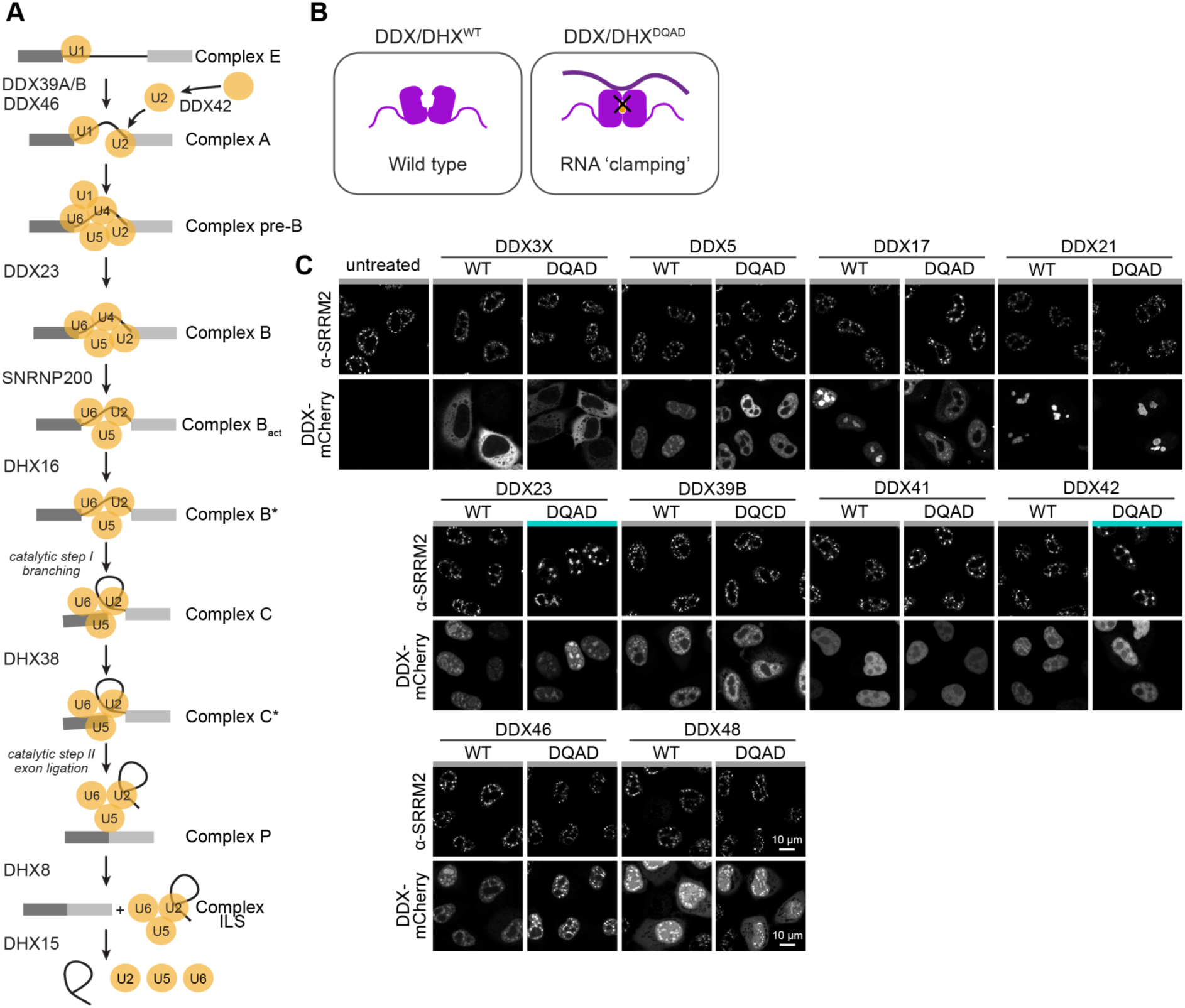
ATPase-deficient DEAD-box helicases selectively remodel nuclear speckle morphology. **(A)** Schematic of the splicing reaction, highlighting the key ATPases that catalyze transitions between successive spliceosomal complexes (E → A → B → B* → C → P → ILS). **(B)** Schematic of the DDX/DHX ATPase mutants used in this study. Wild-type (WT) helicases engage RNA transiently, coupling ATP binding and hydrolysis to substrate release and thereby driving directional mRNA flux through the spliceosome. Substitution of the catalytic Walker-B motif (DEAD→DQAD in DDX, DEAH→AQAD in DHX) abolishes ATP hydrolysis, trapping the enzyme on RNA in a high-affinity, non-productive state (“RNA clamping”). **(C)** HeLa Kyoto cells were transiently transfected to express C-terminally mCherry-tagged DEAD-box (DDX) ATPases, wild-type (WT) or ATPase-deficient (DQAD) mutants. 24 h after transfection, cells were fixed and immunostained for SRRM2 using the SC35 antibody. Representative maximum-intensity projections of Z-stacks are shown. Turquoise color bars mark conditions with enlarged nuclear speckles. N = 3, n ≥ 60. Scale bars, 10 µm.

**Figure S2.**
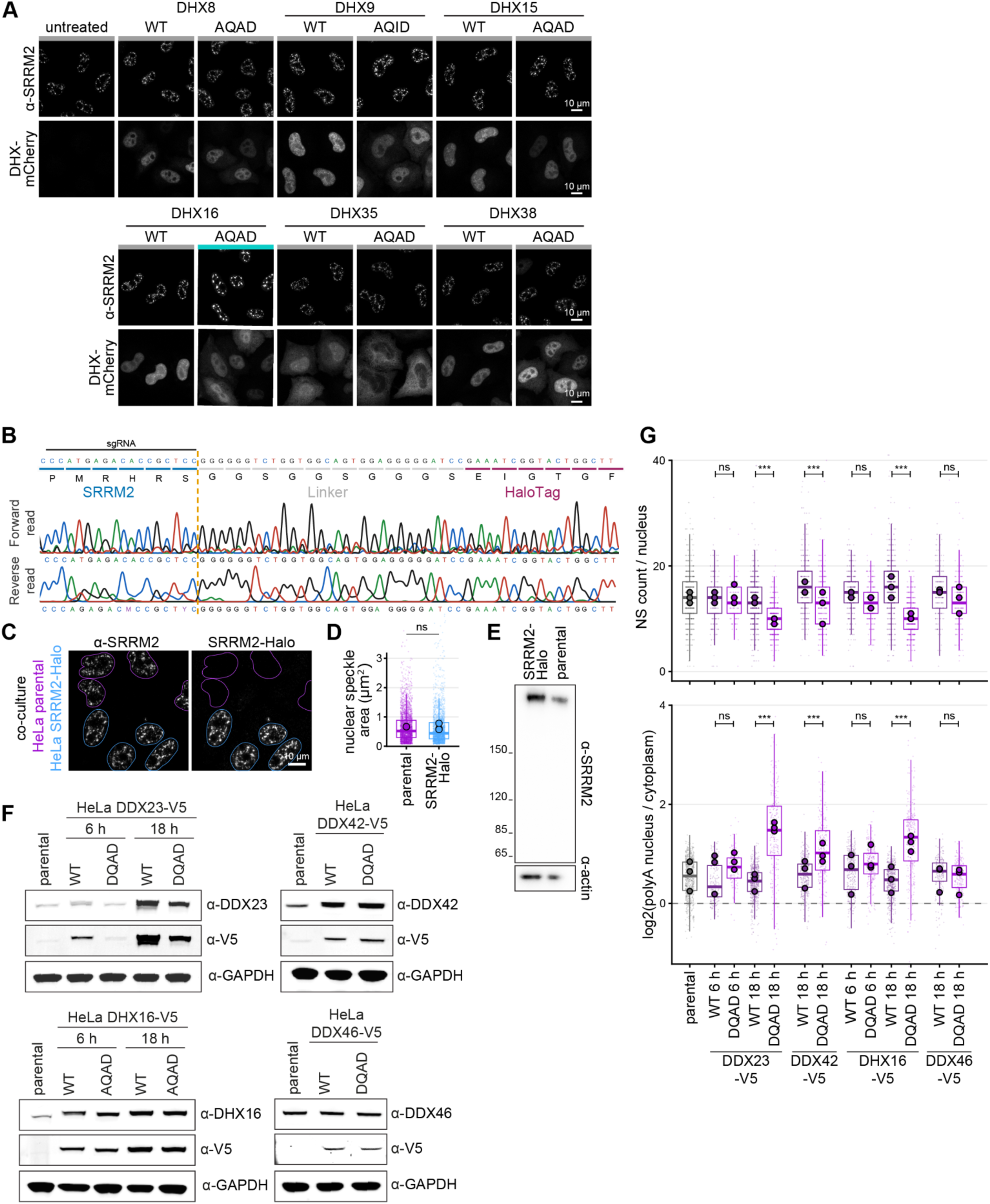
ATPase-deficient DExH-box helicases remodel nuclear speckles and promote nuclear poly(A)+ RNA accumulation. **(A)** HeLa Kyoto cells were transiently transfected to express C-terminally mCherry-tagged DExH-box (DHX) ATPases, wild-type (WT) or ATPase-deficient (AQAD) mutants. 24 h after transfection, cells were fixed and immunostained for SRRM2 using the SC35 antibody. Representative maximum-intensity projections of Z-stacks are shown. Turquoise color bars mark conditions with enlarged nuclear speckles. N = 3, n ≥ 68. Scale bars, 10 µm. **(B)** Validation of the endogenous SRRM2-HaloTag knock-in generated by CRISPaint. A HaloTag cassette was inserted at the SRRM2 C-terminus using the sgRNA 5′- CCATGAGACACCGCTCCTCC-3′, producing an in-frame fusion via a (Gly-Gly-Ser)₂-Gly- Gly-Gly-Ser linker. The junction was confirmed by Sanger sequencing. **(C)** Parental HeLa cells and HeLa SRRM2-Halo knock-in cells were co-cultured, cells were treated with TMR-Halo dye for 18 h before fixation and immunostained against SRRM2. Representative maximum-intensity projections are shown. N = 3. n ≥ 104. Scale bars, 10 µm. **(D)** Quantification of nuclear speckle area of co-cultured cells in (D). Cell lines were distinguished post-hoc on a per-nucleus basis by HaloTag (TMR) mean intensity. In each plot, small semi-transparent points are individual measurements; box plots show the median and interquartile range of the pooled data; and large outlined points are the per-replicate means. N = 3. n(nuclei) ≥ 104, n(nuclear speckles) ≥ 2124. Statistics: comparison of mean of three biological replicates using paired two-tailed t-test, p= 0.84. **(E)** Western blot of Hela SRRM2-Halo knockin cells and parental HeLa cells, immunostained with the indicated antibodies. **(F)** Stable HeLa cell lines were induced with doxycycline to express the selected C- terminally V5-tagged DDX/DHX mutants shown in Figure 1E. Expression of DDX23 and DHX16 mutants were induced for 6 h or 18 h, DDX42 and DDX46 were induced for 18 h. Protein expression was analyzed by fluorescence Western blotting using the indicated antibodies. **(G)** Quantification of induced V5-DDX/DHX cell lines in Figure 1E. Number of nuclear speckles per nucleus and nuclear-to-cytoplasmic poly(A) ratio are shown. Small points represent individual cells, boxplots show the cell-level distributions, and large circles indicate the mean of each biological replicate. Three biological replicates were analyzed. Statistical significance was assessed by comparing indicated matched WT–mutant using hierarchical linear mixed-effects models accounting for biological replicate, condition within replicate and imaging field, followed by planned contrasts with Benjamini–Hochberg correction across the six comparisons. For count: p(DDX23 WT vs DQAD, 18h)= 6.84×10^−5^; p(DDX42 WT vs DQAD, 18h)= 1.01×10^−4^; p(DHX16 WT vs AQAD, 18h)=1.01×10^−10^. For nuclear-to-cytoplasmic poly(A) ratios: p(DDX23 WT vs DQAD, 18h)= 2.46×10^−12^; p(DDX42 WT vs DQAD, 18h)= 7.9×10^−4^; p(DHX16 WT vs AQAD, 18h)=1.35×10^−8^.

**Figure S3:**
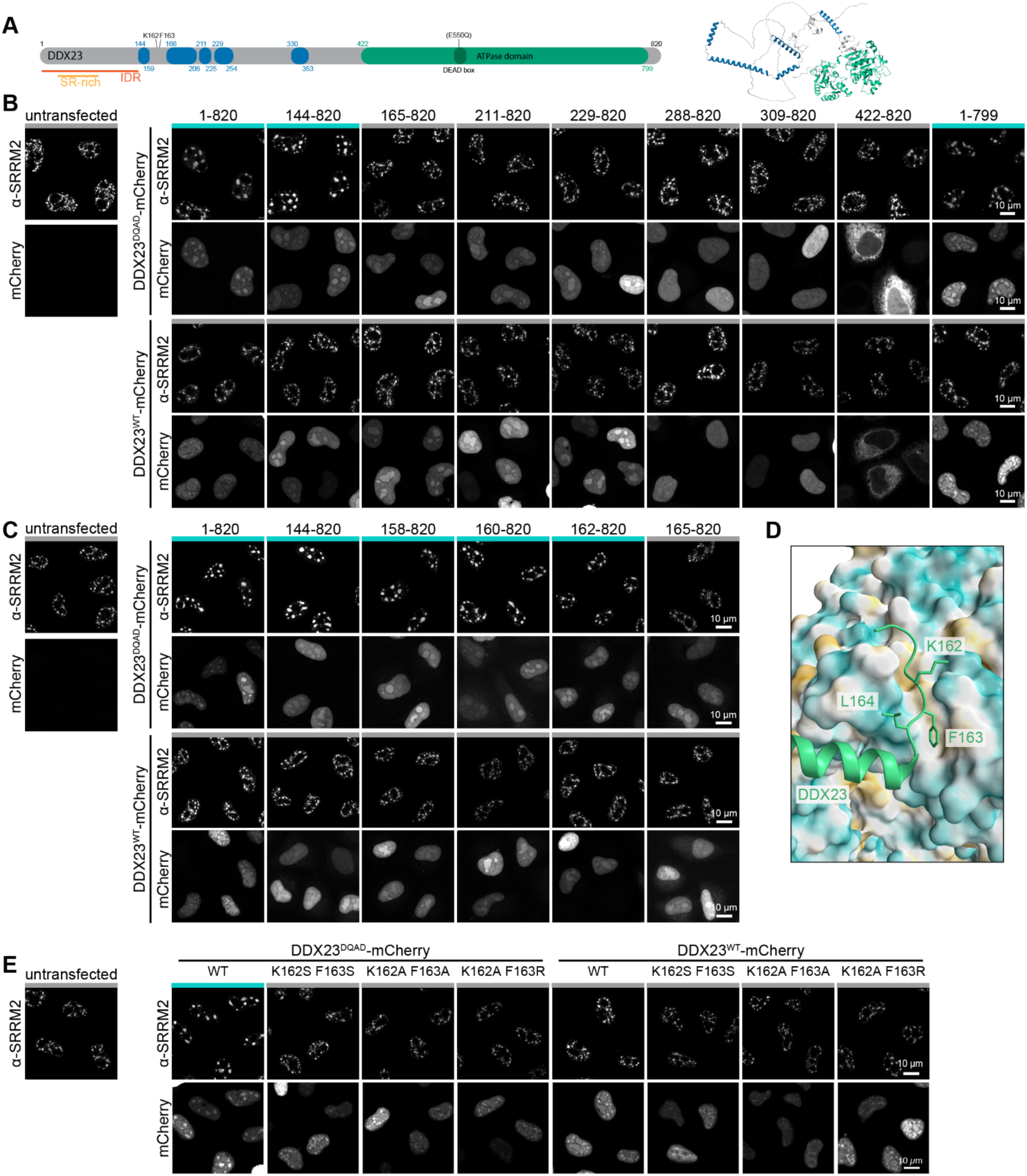
A DDX23 N-terminal SNRNP200-interaction motif is required for DDX23-DQAD-induced nuclear speckle remodeling. **(A)** Domain organization of DDX23 (816 aa): the C-terminal ATPase domain (green, ∼422– 799) and N-terminal predicted α-helices and intervening unstructured regions (blue, residue boundaries indicated), alongside the AlphaFold model colored by the same scheme. Orange bars below the schematic mark the N-terminal intrinsically disordered region (IDR, residues 1–145; AlphaFold pLDDT < 70) and, within it, the single SR-rich block (residues 21–74; >45% Ser+Arg). **(B)** Coarse truncation series of DDX23. HeLa Kyoto cells were transiently transfected with the indicated C-terminally mCherry-tagged DDX23 truncations, in the WT or ATPase-deficient (DQAD) background. Cells were fixed 24 h after transfection and immunostained for SRRM2 using the SC35 antibody. Representative maximum-intensity projections of Z-stacks are shown. Turquoise color bars mark conditions with enlarged nuclear speckles. N = 3. Scale bars, 10 µm. **(C)** Fine truncation series narrowing the N-terminal boundary (residues ∼144–165), otherwise as in (B). **(D)** Structural context of the identified DDX23 N-terminal motif at its interface with SNRNP200. The SNRNP200 surface is colored by hydrophobicity (hydrophobic, gold; hydrophilic, cyan); DDX23 is shown in green with residues K162, F163 and L164 highlighted. PDB: 6QW6. **(E)** DDX23 point-mutant analysis. HeLa Kyoto cells were transiently transfected with C-terminally mCherry-tagged DDX23 (WT or DQAD) carrying the indicated substitutions at K162/F163 (K162S F163S, K162A F163A, K162A F163R). Fixation, staining and imaging as in (B). N = 3. Scale bars, 10 µm.

**Figure S4:**
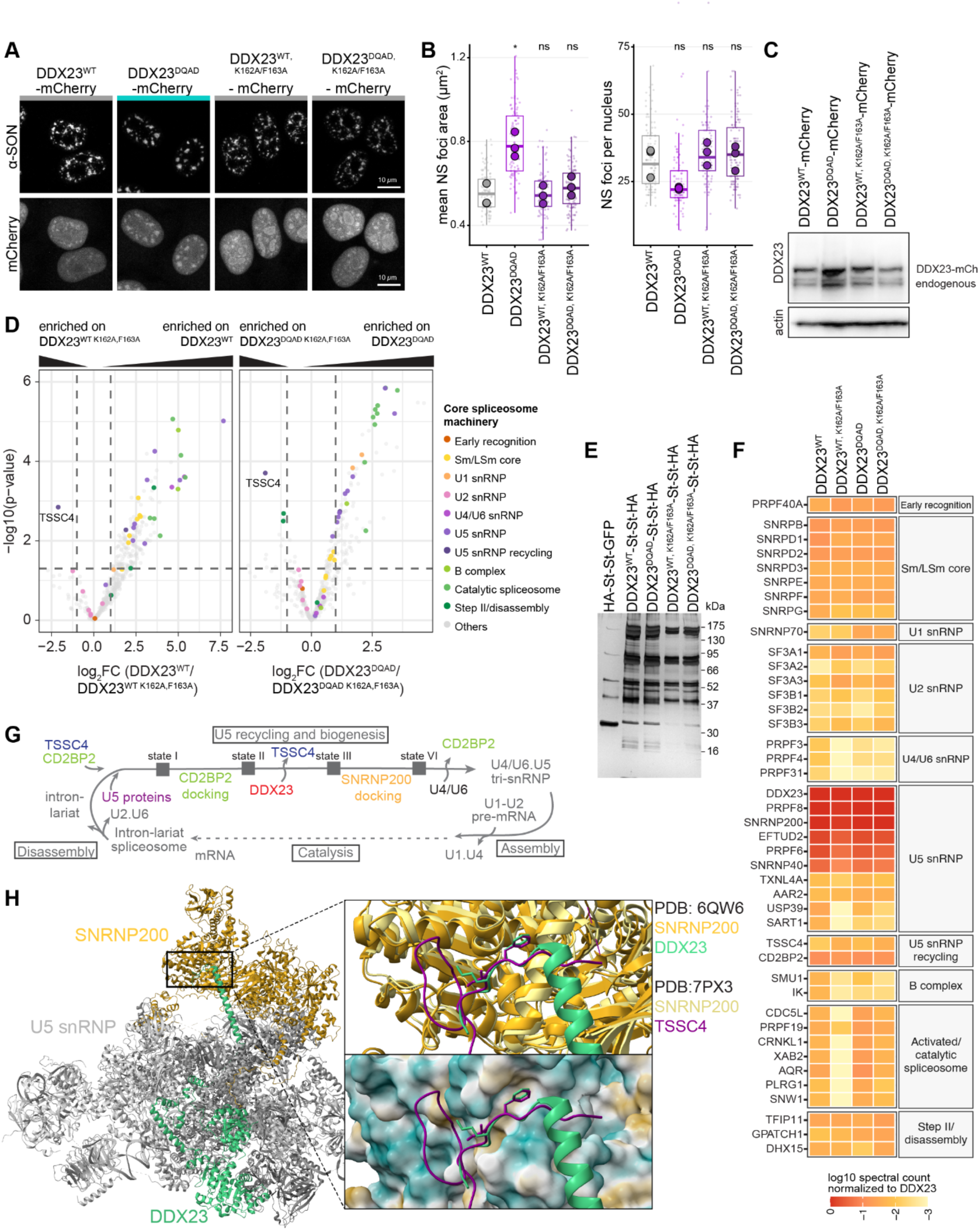
Disruption of the DDX23–SNRNP200 interface traps a TSSC4-associated U5 snRNP state. **(A)** Stable HeLa cell lines were induced with doxycycline to express C-terminally mCherry-tagged DDX23 variants (WT, DQAD, WT^K162A/F163A^, DQAD^K162A/F163A^) at near-endogenous levels for 24 h. After fixation, cells were immunostained for SON. Representative maximum-intensity projections are shown. Turquoise color bars mark conditions with enlarged nuclear speckles. N = 3, n ≥ 97. Scale bars, 10 µm. **(B)** Quantification of stable cell lines expressing DDX23 mutants in (A). Nuclear speckle number per nucleus shown as boxplots. Small points represent individual cells, boxplots show the cell-level distributions, and large circles indicate the median of each biological replicate. Statistical significance was assessed against DDX23 wildtype using a two-tailed paired t-test on biological replicate means. p(DDX23^DQAD^ vs DDX23^WT^) = 0.015; ns, not significant. **(C)** Stable HeLa cell lines induced with doxycycline to express C-terminally mCherry-tagged DDX23 variants in (A), were analyzed for protein expression by Western blot with the indicated antibodies. **(D)** Affinity purification–mass spectrometry (AP-MS) of DDX23 variants. Stable HEK293T cell lines were induced to express C-terminally TwinStrep-HA–tagged DDX23 (WT, WT K162A/F163A, DQAD, or DQAD K162A/F163A) for 8 h; complexes were purified on Strep-Tactin beads and eluted with biotin. Volcano plots show enrichment of co-purifying proteins versus significance; dashed lines mark thresholds (|log₂ FC| = 1; p = 0.05). Points are colored by spliceosome sub-complex (legend); TSSC4 is labeled. N = 3. **(E)** Silver-stained SDS-PAGE of the eluates from (D). **(F)** Heatmap of normalized spectral counts for splicing factors co-purifying with each DDX23 variant, grouped by functional stage (early recognition, Sm/LSm core, U1/U2/U4-U6/U5 snRNP, U5 snRNP recycling, B complex, activated/catalytic spliceosome, step II/disassembly). Color encodes log₁₀ spectral count normalized to DDX23. **(G)** Schematic of the U5 snRNP recycling and biogenesis pathway. Following spliceosome disassembly, U5 proteins and CD2BP2 assemble the post-catalytic U5 snRNP; CD2BP2 docking, DDX23 recruitment and SNRNP200 docking (states I–VI) regenerate the mature U4/U6.U5 tri-snRNP for a new round of assembly and catalysis. **(H)** Structural positioning of the K162/F163 motif within the U5 snRNP. The DDX23– SNRNP200 complex (PDB 6QW6; SNRNP200 gold, DDX23 green) is overlaid with SNRNP200 bound to TSSC4 (PDB 7PX3; TSSC4 purple). The inset shows that phenylalanine F215 of TSSC4 and F163 of DDX23 occupy the same hydrophobic pocket on SNRNP200 (surface colored by hydrophobicity; hydrophobic, gold; hydrophilic, cyan).

**Figure S5:**
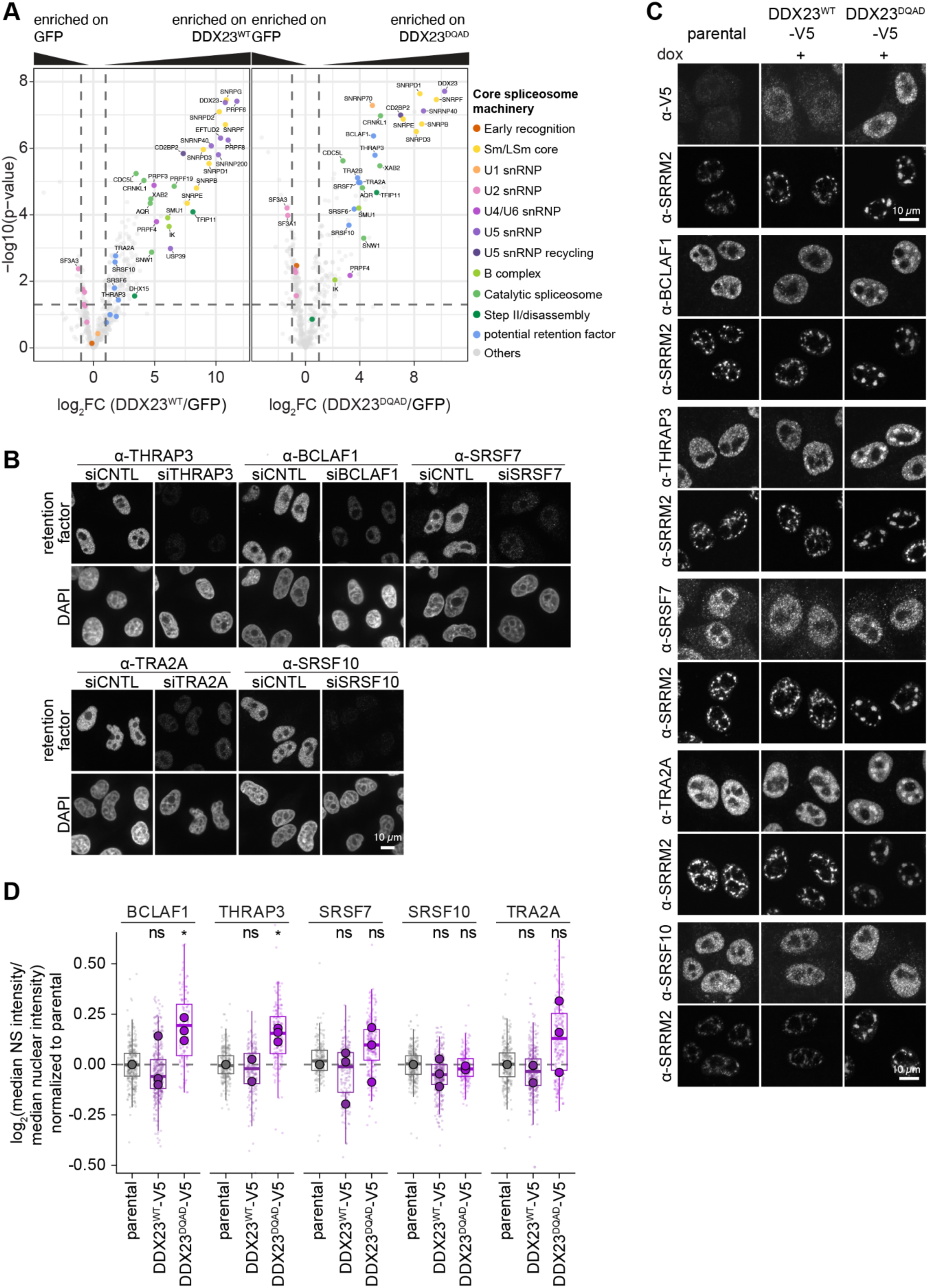
Candidate retention factors selectively associate with DDX23-DQAD complexes and accumulate in nuclear speckles. **(A)** Volcano plots of the DDX23 AP-MS dataset (related to Fig 2A and Fig S4D), showing enrichment of co-purifying proteins on DDX23^WT^-Strep-Strep-HA or DDX23^DQAD^-Strep-Strep-HA relative to a HA-Strep-Strep-GFP control; dashed lines mark thresholds (|log₂ FC| = 1 and p = 0.05). Points are colored by spliceosome sub-complex and potential retention factors. N = 3. **(B)** Antibody validation. HeLa cells were depleted of the indicated proteins by siRNA or treated with control siRNA (siCNTL), then immunostained with the corresponding antibody to confirm antibody signal specificity. **(C)** Stable HeLa cell lines were induced with doxycycline to express DDX23^WT^ or DDX23^DQAD^ (V5-tagged) at near-endogenous levels for 24 h. After fixation, cells were immunostained with the indicated proteins. Representative maximum-intensity projections are shown. N = 3, n ≥ 128. Scale bars, 10 µm. **(D)** Quantification of (C). Median speckle/whole-nucleus intensity ratio per cell for potential pre-mRNP retention factors (BCLAF1, THRAP3, SRSF7, SRSF10, TRA2A), normalized within each replicate to the SRRM2-Halo parental control and log₂-transformed (0 = parental, dashed line). Boxes, median and IQR of single cells (faint points); large circles, per-replicate means. Significance was tested on replicate means by two-tailed paired t-test versus the matched SRRM2-Halo parental. DDX23-DQAD 24 h increased speckle enrichment 1.13-fold for BCLAF1 (p = 0.034) and 1.11-fold for THRAP3 (p = 0.017); all other DDX23-DQAD and all DDX23-WT comparisons were not significant (p > 0.05).

**Figure S6:**
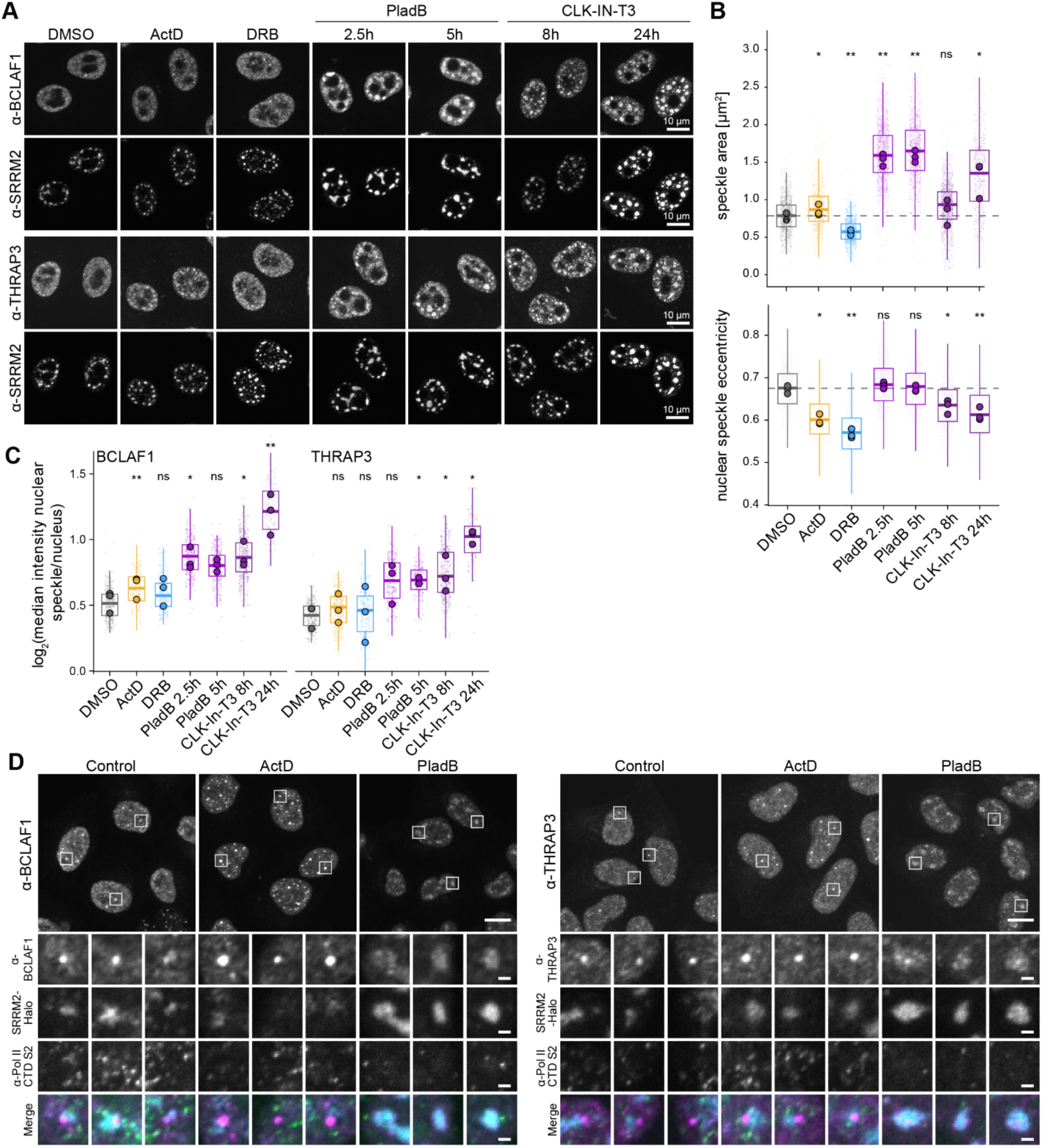
Splicing and transcriptional perturbations differentially remodel nuclear speckles and retention-factor localization. **(A)** HeLa cells were treated with DMSO, actinomycin D (ActD,1 µg/mL, 2.5 h), DRB (50 µg/mL, 2.5 h), pladienolide B (PladB, 500 nM, 2.5 h or 5 h), or the CLK inhibitor CLK-IN-T3 (1 µM, 8 h or 24 h), as indicated. After fixation, cells were immunostained with the indicated antibodies. Representative maximum-intensity projections are shown. N = 3, n ≥ 83 cells per condition. Scale bars, 10 µm. **(B)** Quantification of nuclear speckle area and nuclear speckle eccentricity from the images in (A), pooled across stainings. Each small dot is a single nucleus; large dots are the per-replicate means (N = 3); boxes show the median and interquartile range of the single-cell distribution; the dashed line marks the DMSO median. Splicing inhibition enlarged speckles, whereas transcription inhibition had opposing effects: relative to DMSO (0.75 µm²), speckle area increased to 1.54 µm² with PladB 2.5 h (p = 0.0053), 1.58 µm² with PladB 5 h (p = 0.0089) and 1.23 µm² with CLK-IN-T3 24 h (p = 0.031), and decreased to 0.56 µm² with DRB (p = 0.0058); ActD caused a smaller increase to 0.85 µm² (p = 0.016). CLK-IN-T3 8 h was not significant (0.85 µm², p = 0.45). Paired two-tailed t-tests of the three replicate means, each treatment against the replicate-matched DMSO control (uncorrected). n ≥ 529 cells per condition. *p < 0.05, **p < 0.01, ns = not significant. **(C)** Quantification of nuclear speckle enrichment of BCLAF1 (left) and THRAP3 (right) from the images in (A), expressed as log₂ of the ratio of median antibody signal inside speckles to the median signal over the whole nucleus (0 = no enrichment). Each small dot is a single nucleus; large dots are the per-replicate means (N = 3); boxes show the median and interquartile range of the single-cell distribution. Statistical significance was assessed on the three replicate means by paired two-tailed t-tests of each treatment against the replicate-matched DMSO control (uncorrected). pValues: p(BCLAF1+ ActD) = 0.0012, p(BCLAF1+PladB 2.5h = 0.021), p(BCLAF1+CLK-IN-T3 8 h) = 0.021, p(BCLAF1+CLK-In-T3)= 0.0049, p(THRAP3+PladB 5 h) = 0.028, p(THRAP3+CLK-IN-T3 8 h)= 0.027 and p(THRAP3+ CLK-IN-T3 24 h) = 0.014. *p < 0.05, **p < 0.01, ns = not significant. **(D)** HeLa SRRM2-Halo cells were labelled with TMR-Halo dye for 18h and treated with ActinomycinD (ActD, 1 μg/mL, 2.5h), or PladienolideB (PladB, 500 nM, 2.5h). After fixation, cells were stained with the indicated antibodies. Representative maximum-intensity projections are shown in the overview panel, enlarged views of single planes are shown below. Merge: BCLAF1/THRAP3 (magenta), SRRM2-Halo (cyan), Poll II CTD S2 (green). N = 3, Scale bars, 10 µm; enlargements, 1 µm.

**Figure S7:**
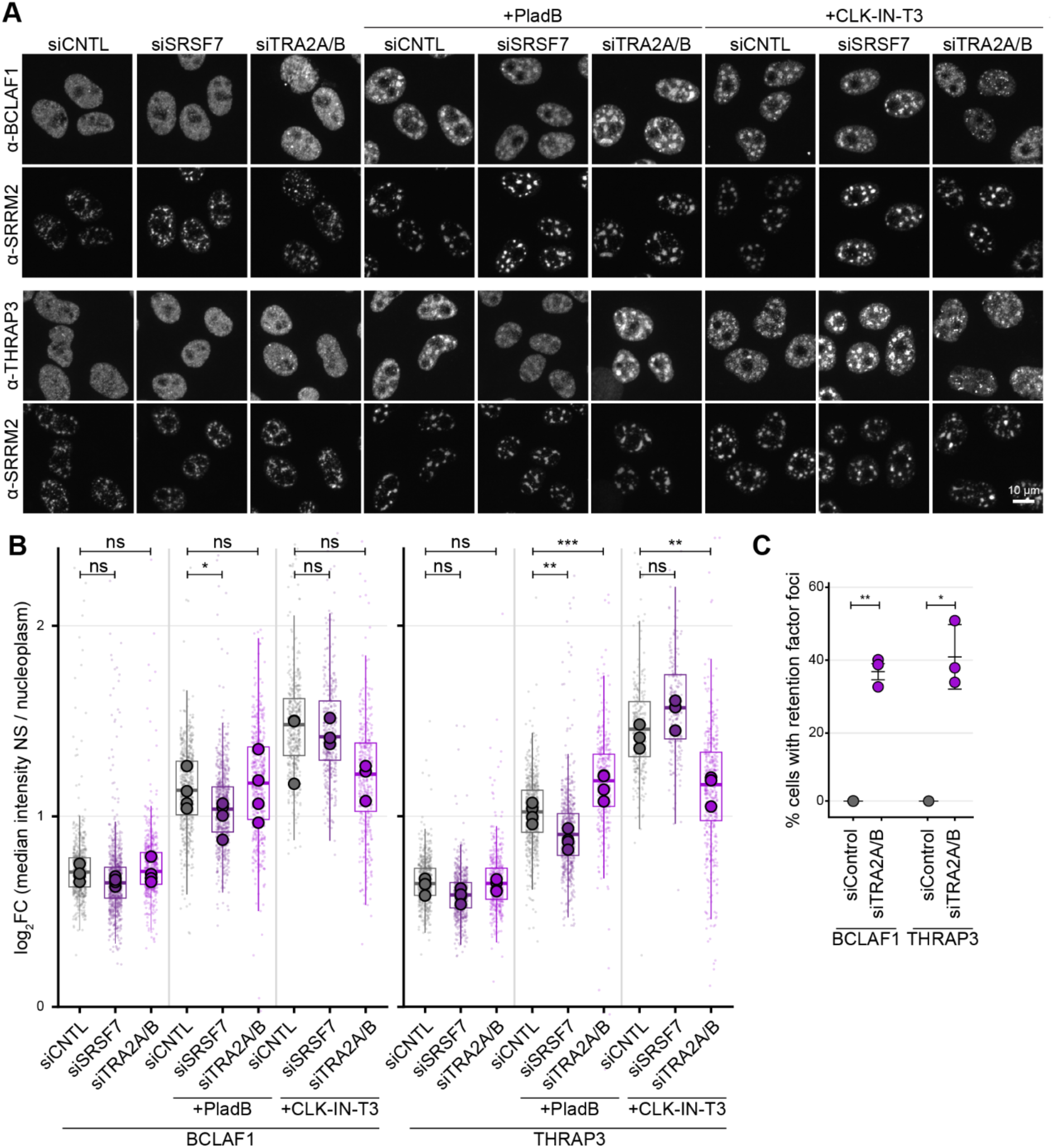
Retention-factor recruitment shows distinct dependencies on SRSF7 and TRA2A/B. **(A)** HeLa cells were depleted of the indicated proteins by siRNA, before fixation, they were either left untreated or treated with pladienolide B (PladB, 500 nM, 2.5 h) or CLK-In-T3 (1 µM, 24 h), and immunostained with the indicated antibodies. Representative maximum-intensity projections are shown. N = 3, n ≥ 229. Scale bars, 10 µm. **(B)** Quantification of (A). For each cell, retention factor enrichment was calculated as log₂ of the ratio of the median retention factor intensity within SRRM2-defined speckles to the median retention factor intensity across the nucleoplasm; a value of 0 indicates no speckle enrichment. Box plots show the distribution across pooled single cells (line, median; box, interquartile range; whiskers, 1.5× IQR; outliers not shown); small points are individual cells and large points are the medians of the biological replicates (colour, siRNA). Statistical significance was assessed by a two-tailed paired t-test on the per-replicate median log₂FC, comparing each knockdown to the matched siCNTL within each treatment. N = 4 for untreated and PladB treated conditions, N=3 for CLK-In-T3 conditions. n ≥ 229 cells per condition. p(THRAP3, PladB, siSRSF7)= 0.0051, p(THRAP3, PladB, siTRA2A/B)= 0.001, p(THRAP2, CLK-In-T3, siTRA2A/B)= 0.0077, p(BCLAF1, PladB, siSRSF7)= 0.0433, ns, not significant. **(C)** Quantification of (A). Percentage of cells displaying THRAP3 or BCLAF1 foci discrete from nuclear speckles. N = 3, n ≥ 340. Statistical significance was assessed two-tailed paired t-test on the per-replicate percentages.

**Figure S8:**
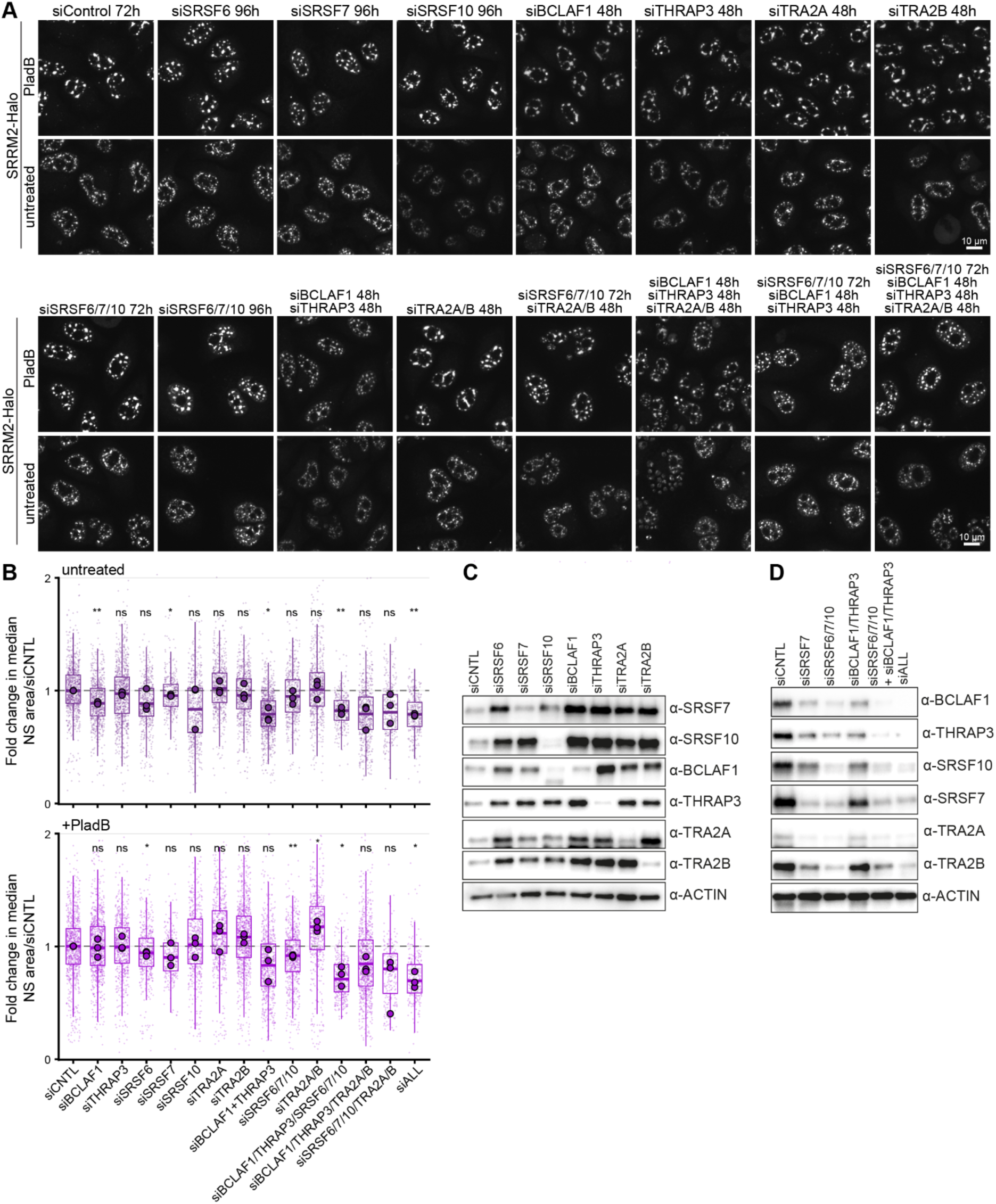
Individual and combined depletion of candidate retention factors differentially alters nuclear speckle morphology. **(A)** HeLa SRRM2-Halo cells were depleted of the indicated factors by siRNA, individually (top block) or in the indicated combinations (bottom block), for the times shown. Cells were treated with TMR-Halo dye for 18 h and either left untreated or treated with pladienolide B (PladB, 500 nM, 2.5 h) before fixation. Representative maximum-intensity projections are shown. N = 3, n ≥ 251. Scale bars, 10 µm. **(B)** Quantification of nuclear speckle area in (A). Mean nuclear speckle area per nucleus was normalized within each biological replicate and treatment to the median of the corresponding siCNTL condition. Small points represent individual cells, boxplots show the cell-level distributions, and large circles indicate the median of each biological replicate. Three biological replicates were analyzed. Statistical significance was assessed for each depletion relative to the treatment-matched siCNTL by one-sample t-test of the replicate-level median fold changes against 1. For untreated: p(siBCLAF1) = 0.00316; p(siSRSF7) = 0.0155; p(siBCLAF1+THRAP3) = 0.0251; p(siBCLAF1/THRAP3/SRSF6/7/10) = 0.00930; p(siALL) = 0.00100. For +PladB: p(siSRSF6) = 0.0327; p(siSRSF6/7/10) = 0.00452; p(siTRA2A/B) = 0.0218; p(siBCLAF1/THRAP3/SRSF6/7/10) = 0.0348; p(siALL) = 0.0178. **(C)** Hela cells depleted of individual proteins by RNAi in (A) were analyzed by Western blot and immunostained with the indicated antibodies. **(D)** Hela cells depleted of individual proteins and protein combinations by RNAi in Fig 2D were analyzed by Western blot and immunostained with the indicated antibodies.

**Figure S9:**
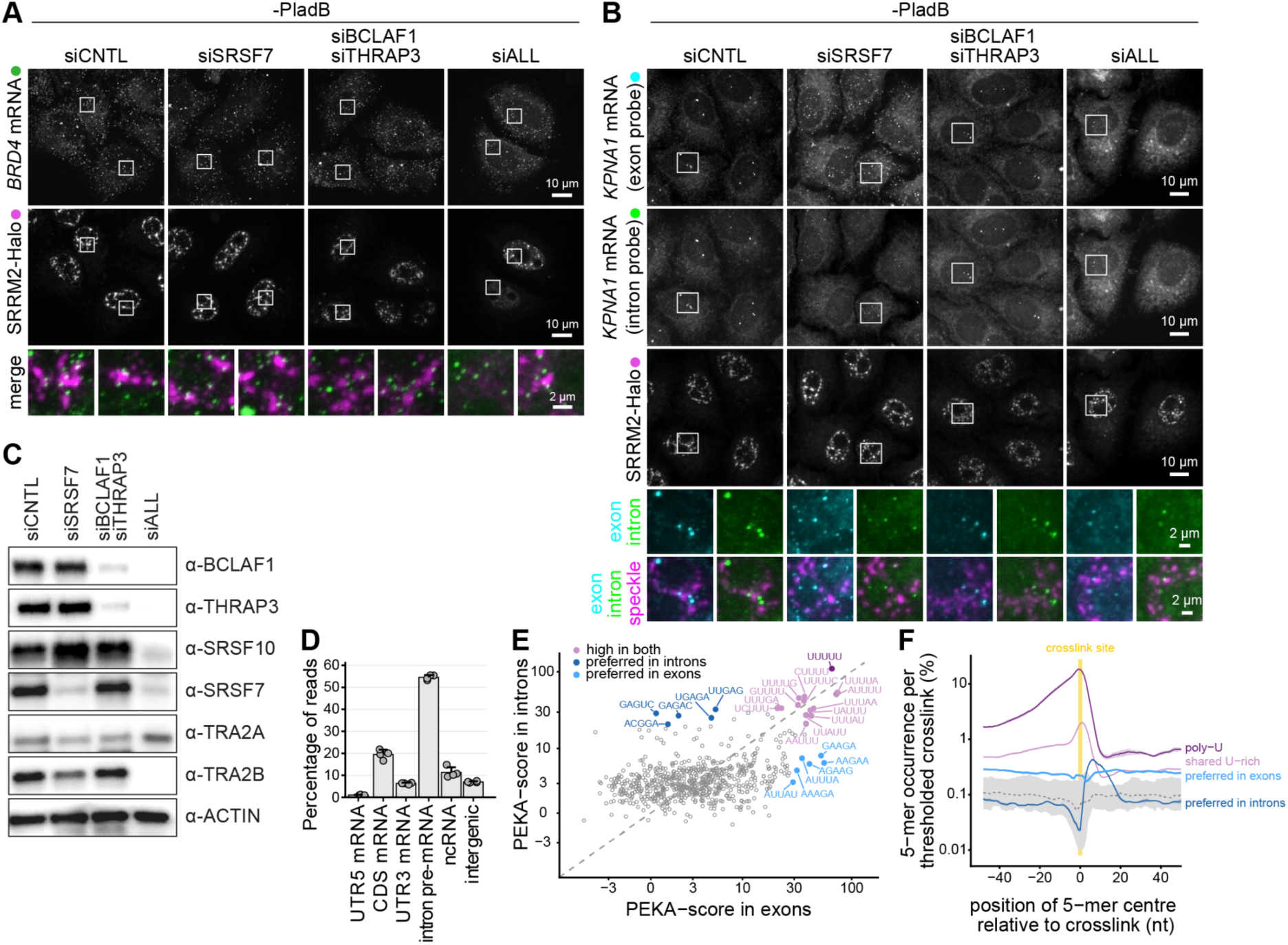
Retention-factor depletion alters the association of BRD4 and KPNA1 mRNAs with nuclear speckles in untreated cells. **(A)** HeLa SRRM2–Halo knockin cells were depleted of the indicated factors by siRNA and labelled with Janelia Fluor 503 HaloTag ligand for 18 h (same experiment as Figure 3A/B). Cells were left untreated (−PladB), fixed, and BRD4 mRNA was detected by smFISH using probes targeting exonic sequences. Representative maximum-intensity projections are shown together with enlarged merged views of the boxed regions. N = 3 biological replicates, n ≥ 209 cells per condition. Scale bars, 10 µm; enlargements, 2 µm. **(B)** HeLa SRRM2–Halo knockin cells were depleted of the indicated factors by siRNA and labelled with Janelia Fluor 503 HaloTag ligand for 18 h (same experiment as Figure 3C/D). Cells were left untreated (−PladB), fixed, and KPNA1 mRNA was detected by smFISH using probe sets targeting either exonic or intronic sequences. Representative maximum-intensity projections are shown together with enlarged views of the boxed regions. N = 3 biological replicates, n ≥ 230 cells per condition. Scale bars, 10 µm; enlargements, 2 µm. **(C)** HeLa cells from Figure 3A-D and Figure S9A/B were analyzed by Western blot using the indicated antibodies. **(D)** Proportion of mapped HA-THRAP3 iCLIP crosslinks falling in different genomic regions. N=4. **(E)** PEKA-score of each 5-mer in introns versus exons, from four THRAP3 iCLIP replicates. Points are the mean of replicates, shown if 5-mers occur in at least three replicates. Exons comprise coding exons and 5′UTRs (PEKA region “other_exon”). Dashed line marks equal enrichment. **(F)** Occurrence of the same groups at each position relative to intronic crosslinks, with the shared group split into UUUUU and the 13 remaining 5-mers. Lines are the mean over group members and samples, smoothed with a centred 5-nt rolling mean as in PEKA’s own cluster plots; coloured shading is the SEM across samples. The grey band is the position-wise 5th– 95th percentile of the 47 5-mers whose mean intronic PEKA-score lies within ±1 of zero, with their median dashed; the shaded column marks the crosslinked nucleotide.

**Figure S10:**
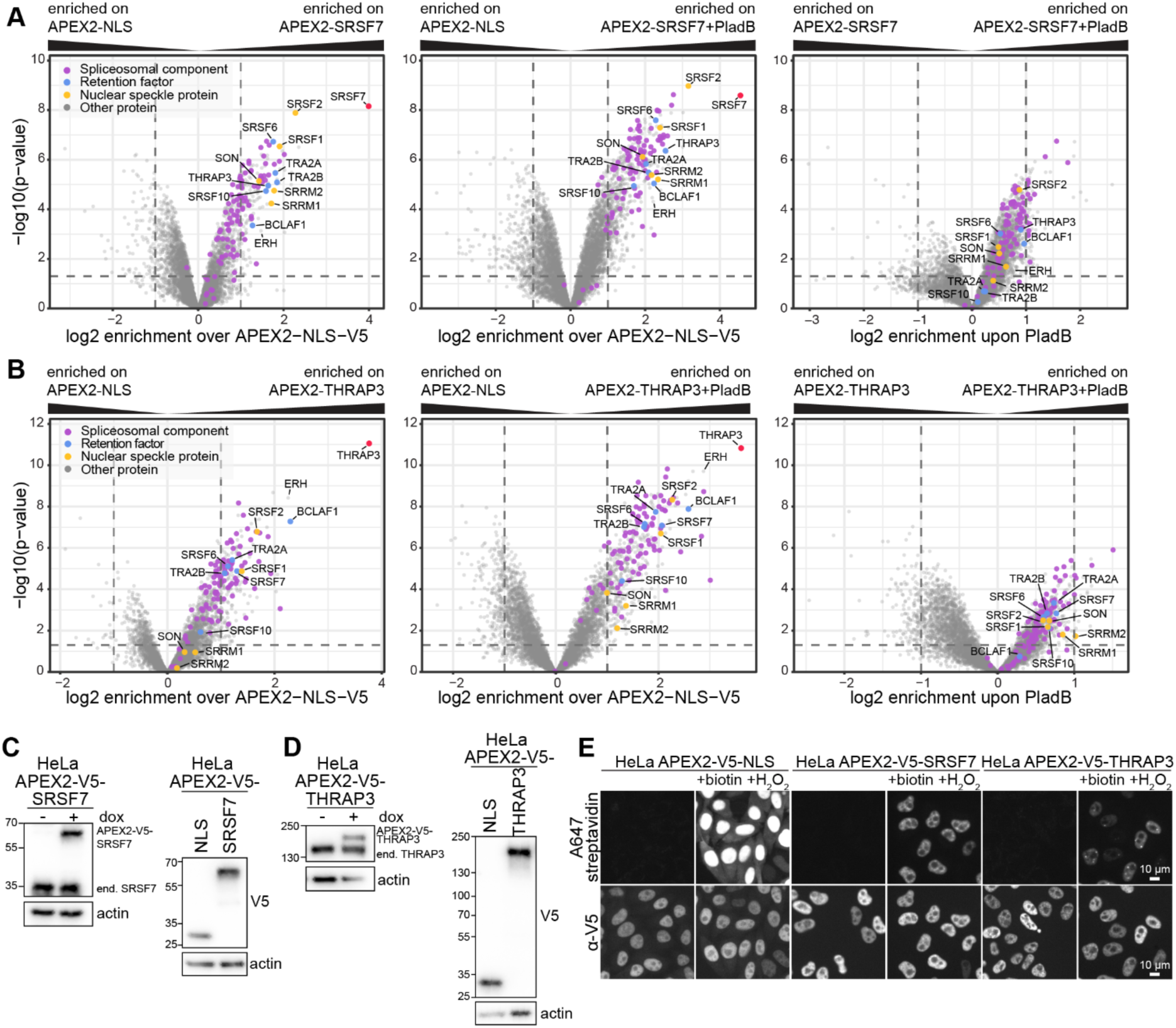
APEX2 proximity proteomics maps the SRSF7 and THRAP3 proximity networks and their response to splicing inhibition. (**A**) APEX2 proximity proteomics of SRSF7. Stable HeLa cells were induced to express APEX2-V5-SRSF7 at near-endogenous levels for 48 h and either left untreated or treated with pladienolide B (PladB, 500 nM, 2.5 h) before APEX2-mediated proximity labeling and harvest; cells expressing APEX2-V5-NLS served as a control. Volcano plots show protein enrichment versus significance, dashed lines mark thresholds (|log₂ FC| = 1; p = 0.05). Points are colored by category (spliceosomal component, retention factor, nuclear speckle protein, other); bait in red. N = 3. (**B**) APEX2 proximity proteomics of THRAP3. Stable HeLa cells were induced to express APEX2-V5-THRAP3 at near-endogenous levels for 48 h and either left untreated or treated with pladienolide B (PladB, 500 nM, 2.5 h) before APEX2-mediated proximity labeling and harvest; cells expressing APEX2-V5-NLS served as a control. Volcano plots show protein enrichment versus significance, dashed lines mark thresholds (|log₂ FC| = 1; p = 0.05). Points are colored by category (spliceosomal component, retention factor, nuclear speckle protein, other); bait in red. N = 3. **(C, D)** Western blots of HeLa APEX2-V5-SRSF7 (C) and APEX2-V5-THRAP3 (D) lines ± doxycycline induced as in (A) and (B), probed with antibodies against bait and actin (loading control). V5 blots (right) confirm expression of the APEX2-V5-NLS control and the respective bait. **(E)** Validation of APEX2 activity and localization. HeLa APEX2-V5-NLS, -SRSF7 and - THRAP3 cells were induced as in (A) and (B), labeled ± biotin-phenol and ± H₂O₂ and imaged for biotinylation (A647-streptavidin) and bait expression (V5). Scale bars, 10 µm.

**Figure S11:**
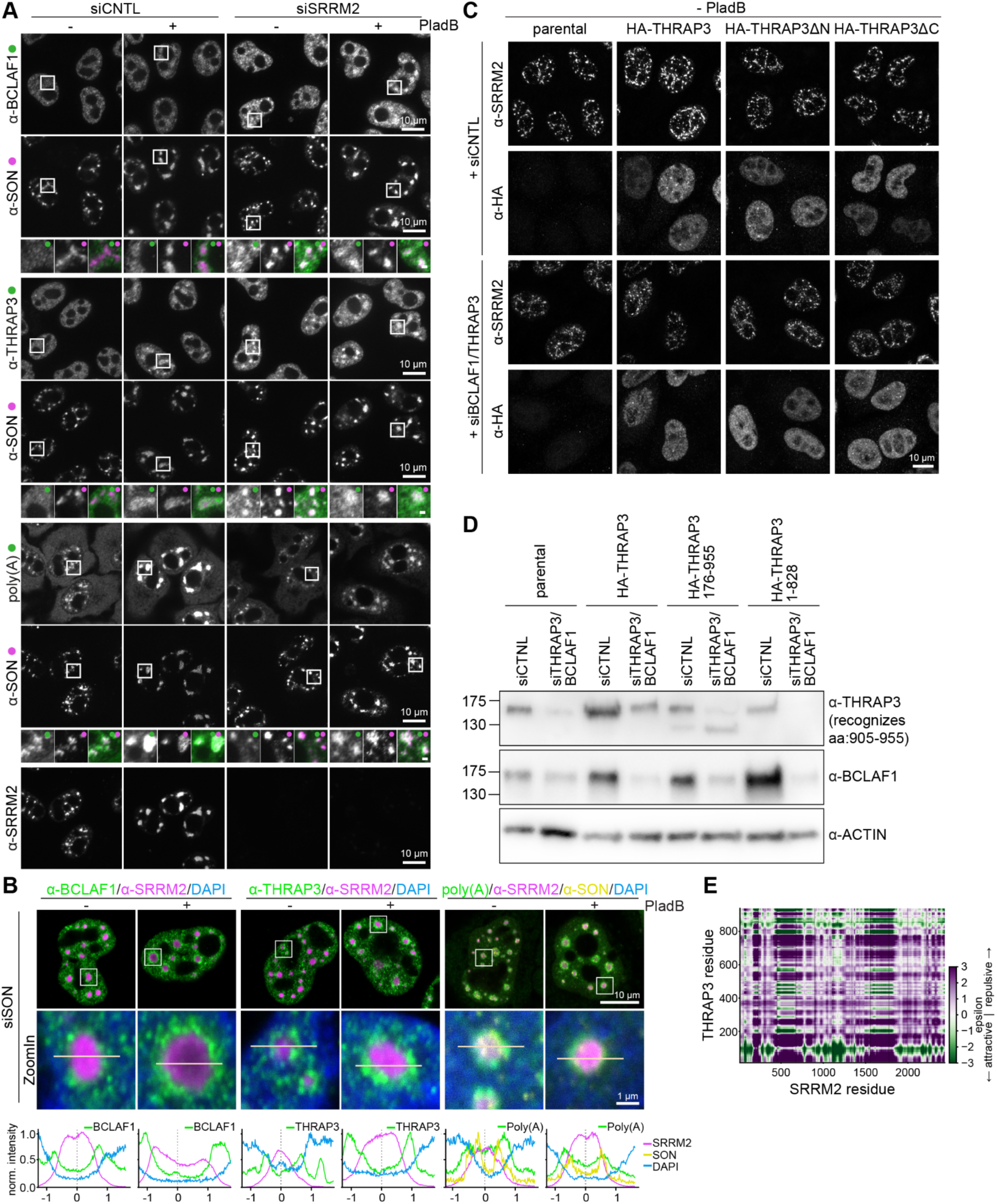
SON and SRRM2 differentially organize retention factors and poly(A)+ RNA within nuclear speckles. **(A)** HeLa were depleted of SRRM2 by siRNA (72 h). Cells were then treated with DMSO or PladB (500 nM) for 2.5 h before fixation. Poly(A) RNA was stained using FISH, proteins were then immunostained with the indicated antibodies. Representative single planes are shown. N = 3. Scale bars, 10 µm; enlargement 1 µm. **(B)** Cells depleted of SON by siRNA (48 h) and treated with PladB in Figure 4B, were imaged at on a LSM770 line scanning confocal microscope. Enlargements of the boxed regions and normalized fluorescence intensity profiles along the indicated lines are shown below. Scale bars, 10 µm; enlargements, 1 µm. **(C)** Stable HeLa cell lines were induced to express HA-THRAP3, HA-THRAP3ΔN (HA-THRAP3^176–955^) or HA-THRAP3ΔC (HA-THRAP3^1–828^) at near-endogenous protein level (same experiment as Figure 4D, there cells were treated with pladienolide B). Cells were then either co-depleted for BCLAF1 and THRAP3 by siRNA or treated with control siRNA. After fixation, cells were immunostained with the indicated antibodies. Representative maximum-intensity projections are shown. N = 3, n ≥ 80. Scale bars, 10 µm. **(D)** HeLa cells depleted of indicated proteins and expressing different THRAP3 constructs in (**C**) and Figure 4D were analyzed by Western blot using the indicated antibodies. **(E)** Residue-resolved FINCHES interaction maps for THRAP3 × SRRM2. Each pixel is the predicted ε between the local windows centered on the corresponding THRAP3 residue (y-axis) and partner residue (x-axis); green denotes attraction (ε < 0) and purple repulsion (ε > 0), on the shared scale at right (ε ∈ [−3, 3]). SRRM2 was tiled into non-overlapping ∼250-residue fragments (boundaries placed outside predicted folded islands), each run against full-length THRAP3 and stitched into the full-length maps shown.

**Figure S12:**
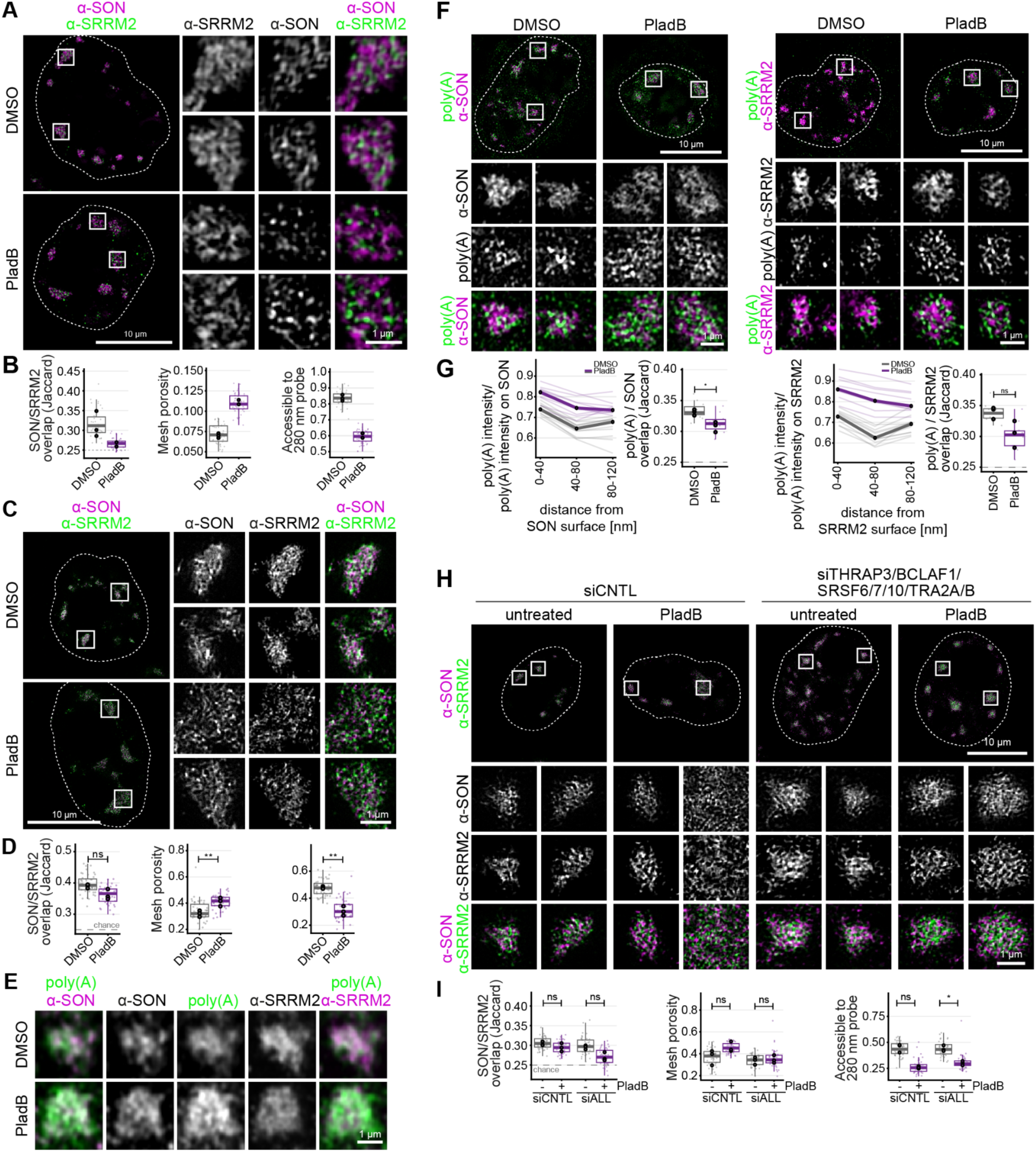
Super-resolution imaging reveals RNA-responsive remodeling of the nuclear speckle scaffold. **(A)** HeLa cells were treated with actinomycin D (1 µg/mL, 2.5 h) or pladienolide B (500 nM, 2.5 h) or DMSO. After fixation, cells were immunostained with anti-SON labelled with a secondary antibody conjugated with Alexa405 and anti-SRRM2 labelled with a secondary antibody conjugated with Alexa488. Reciprocal stainings with swapped secondary colors are displayed in Figure 5A. Cells were imaged using 3D-SIM. Representative reconstructed single planes are shown. N = 2, n ≥ 29. Scale bars, 10 µm, enlargements 1 µm. **(B)** Quantification of cells in (A). Left, overlap of the SON- and SRRM2-dense phases within speckle bodies, measured as an occupancy-matched Jaccard index (top 40% voxels); the dashed line marks the chance level of 0.25, obtained when one channel is rotated 90 degrees within the same nucleus. Middle, mesh porosity of the resolved speckle core (fraction of core volume not occupied by SON/SRRM2 scaffold). Right, fraction of the intra-speckle interstitium reachable by a 280 nm spherical probe entering from the nucleoplasm.Small points, one value per cell (median over that cell’s speckles); large points, per-replicate medians; boxes, cell-level median and interquartile range with whiskers to 1.5x IQR. **(C)** HeLa cells were treated with actinomycin D (1 µg/mL, 2.5 h) or pladienolide B (500 nM, 2.5 h) or DMSO. After fixation, cells were immunostained with the indicated antibodies and imaged using TauSTED. Representative reconstructions of nuclear speckle mid planes are shown. N = 3. Scale bars, 10 µm, enlargements 1 µm. **(D)** Quantification of cells in (C). Left, overlap of the SON- and SRRM2-dense phases within speckle bodies, measured as an occupancy-matched Jaccard index (top 40% voxels); the dashed line marks the chance level of 0.25, obtained when one channel is rotated 90 degrees within the same nucleus. Middle, mesh porosity of the resolved speckle core (fraction of core volume not occupied by SON/SRRM2 scaffold). Right, fraction of the intra-speckle interstitium reachable by a 280 nm spherical probe entering from the nucleoplasm.Small points, one value per cell (median over that cell’s speckles); large points, per-replicate medians; boxes, cell-level median and interquartile range with whiskers to 1.5x IQR. **(E)** HeLa cells were treated with actinomycin D (1 µg/mL, 2.5 h) or pladienolide B (500 nM, 2.5 h) or DMSO. After fixation, poly(A) was stained using FISH, then SON and SRRM2 were immunostained using antibodies. Cells were imaged using LSM880 airyscan. Representative reconstructions of nuclear speckle mid planes are shown. N = 1. Scale bars, 1 µm. **(F)** HeLa cells were treated with pladienolide B (500 nM, 2.5 h) or DMSO. After fixation, poly(A) was stained using FISH (Atto488), then SON or SRRM2 were immunostained using secondary antibodies fused to Alexa405 fluorophores. Cells were imaged using 3D-SIM. Representative reconstructions of nuclear speckle mid planes are shown. N = 3. Scale bars, 10 µm, enlargements 1 µm. **(G)** Quantification of cells in (F). First and third plot: poly(A) intensity inside speckle bodies as a function of distance from the scaffold (SON or SRRM2) surface, expressed relative to the poly(A) intensity on the scaffold of the same cell, so the profile is independent of staining and exposure. Distance bins are 0–40, 40–80 and 80–120. Bins near 40–80 nm include a larger share of the sparse 80 nm boundary layer; the profile is monotonic when that layer is excluded. Thin lines, individual cells; heavy lines, condition medians. Second and forth plots: occupancy-matched overlap of poly(A) with the scaffold (Jaccard of the top 40% of body voxels in each channel; dashed line, chance level 0.25, confirmed empirically by rotating the poly(A) channel 90° in xy: 0.252 DMSO, 0.258 PladB). Points, individual cells; large circles, replicate medians; boxes, median and interquartile range. Statistics: paired two-sided t-test on replicate medians, Holm-corrected across the panels shown; ns, p > 0.05. **(H)** HeLa cells were co-depleted of THRAP3, BCLAF1, SRSF6/7/10, TRA2A/B by siRNA or treated with control siRNA. Before fixation, cells were treated with pladienolide B (500 nM, 2.5 h) or left untreated. After immunostaining with the indicated antibodies, cells were imaged using TauSTED. Representative reconstructed single planes are shown. N = 3. Scale bars, 10 µm, enlargements 1 µm. **(I)** Quantification of cells in (H). Left, overlap of the SON- and SRRM2-dense phases within speckle bodies, measured as an occupancy-matched Jaccard index (top 40% voxels); the dashed line marks the chance level of 0.25, obtained when one channel is rotated 90 degrees within the same nucleus. Middle, mesh porosity of the resolved speckle core (fraction of core volume not occupied by SON/SRRM2 scaffold). Right, fraction of the intra-speckle interstitium reachable by a 280 nm spherical probe entering from the nucleoplasm.Small points, one value per cell (median over that cell’s speckles); large points, per-replicate medians; boxes, cell-level median and interquartile range with whiskers to 1.5x IQR.

